# A non-canonical Arp2/3 complex is essential for *Plasmodium* DNA segregation and transmission of malaria

**DOI:** 10.1101/2023.10.25.563799

**Authors:** Franziska Hentzschel, David Jewanski, Yvonne Sokolowski, Pratika Agarwal, Anna Kraeft, Kolja Hildenbrand, Lilian P. Dorner, Mirko Singer, Friedrich Frischknecht, Matthias Marti

**Affiliations:** Integrative Parasitology, Centre for Infectious Diseases, University Hospital Heidelberg, Heidelberg University Medical Faculty; Heidelberg, Germany; Wellcome Centre for Integrative Parasitology, University of Glasgow; Glasgow, United Kingdom; German Center for Infection Research, DZIF, partner site Heidelberg; Heidelberg, Germany; VetSuisse and Medical Faculties, University of Zurich; Zurich, Switzerland

## Abstract

The malaria-causing parasite *Plasmodium* has a complex life cycle involving both vertebrate and mosquito hosts. Sexual stages or gametocytes are the only stage competent for transmission to mosquitoes. Formation of flagellated male gametes from gametocytes requires rapid rounds of genome replication. Here we discovered a non-canonical *Plasmodium* actin-related protein 2/3 (Arp2/3) complex essential for DNA segregation during male gametogenesis. *Plasmodium* Arp2/3 dynamically localizes within the nucleus to the endomitotic spindles and interacts with a kinetochore protein. Deletion of key Arp2/3 subunits or interfering with actin polymerisation leads to the formation of sub-haploid male gametes and a complete block in transmission through delayed developmental arrest at the oocyst stage. Our work identified an evolutionary divergent protein complex in malaria parasites that offers potential targets for transmission-blocking interventions.

## Introduction

*Plasmodium*, the causative agent of malaria, is an evolutionary distant single-celled apicomplexan parasite. It undergoes a complex life cycle between vertebrate and mosquito hosts where it constantly differentiates into uniquely adapted cellular forms (**Fig. 1A**). While transcription factors governing most of these changes have been identified (*1*), the molecular mechanisms driving most differentiation processes are not well understood. One third of the predicted *Plasmodium* proteins have no homology outside of apicomplexans (*2*). In addition, evolutionary conserved proteins and protein complexes governing fundamental biological processes can be hard to recognize, such as the nuclear porins (*3*), or they are thought to be absent, such as the spindle assembly checkpoint (*4–6*) or the actin nucleator actin-related protein 2/3 (Arp2/3) complex (*7–12*). *Plasmodium* parasites undergo gamete formation and sexual replication in the mosquito. Among the many differentiation processes of *Plasmodium*, male gametogenesis is a particularly complex, fast, and intriguing process, in which eight flagellated microgametes emerge from a parent cell within just 10-15 minutes (*13*, *14*). To achieve this rapid formation of gametes, the cytoplasmic formation of eight axonemes is paralleled in the nucleus by three rounds of closed mitosis without chromosome condensation or nuclear division (endomitosis) (*13*, *15*). Axonemes and mitotic spindles are connected by a bipartite microtubule organising centre that spans the nuclear envelope and contains a basal body on the cytosol and a spindle pole body like structure in the nucleus (*13*, *16*). Once axonemes gain motility, the eight gametes swim out of the parent cell, each pulling a genome along in a process that is both error-prone and poorly understood (*14*). Nuclear division is thought to occur only during gamete budding, and faithful genome segregation from the octoploid nucleus is thus essential for the formation of fertile gametes. However, in absence of a known canonical spindle assembly checkpoint (*5*, *6*, *17*), it is unclear how the parasite safeguards correct chromosome segregation during the rapid rounds of endomitosis and gamete formation.

**Figure 1:**
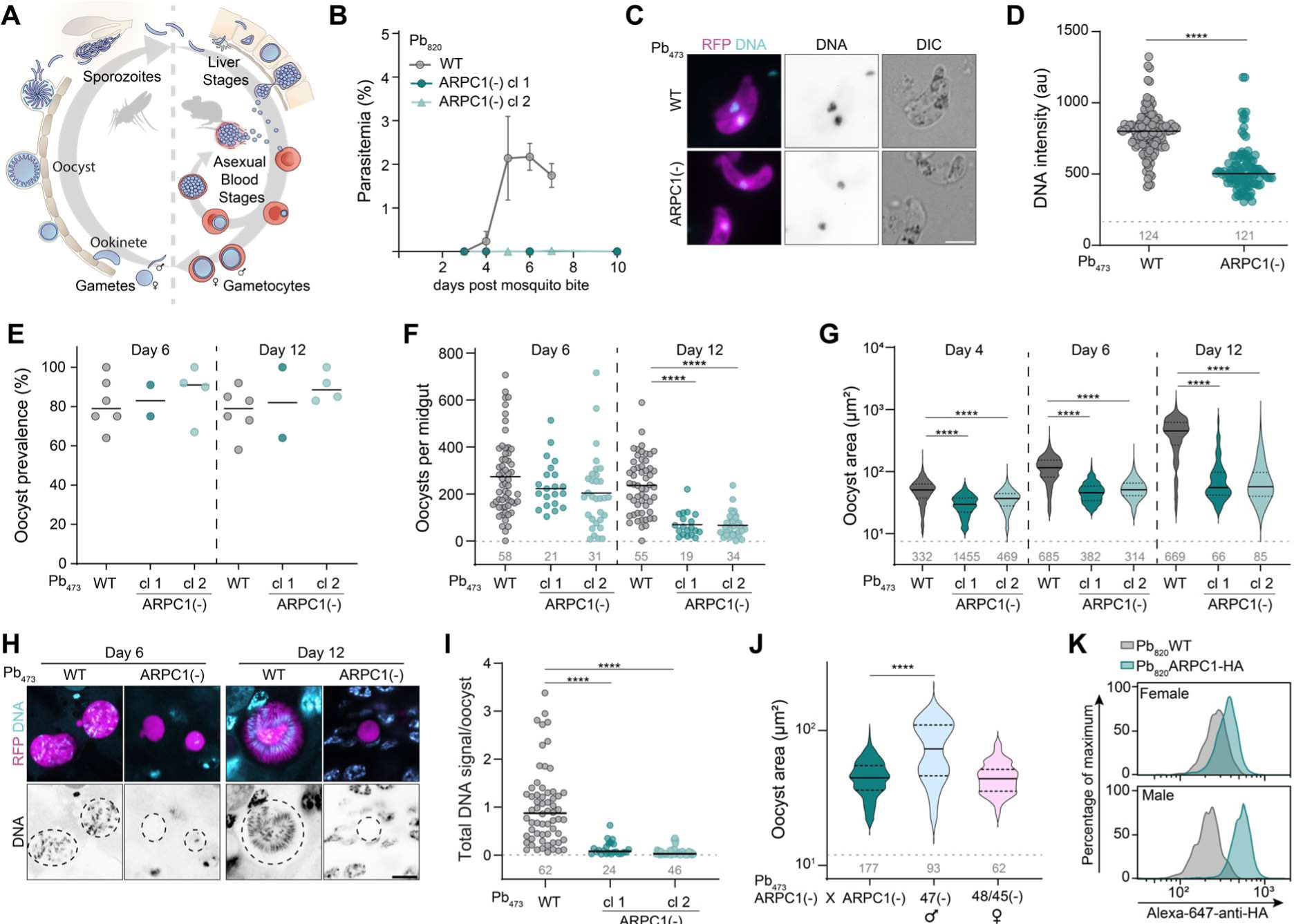
**ARPC1 is essential for parasite transmission. A**) Life cycle of *Plasmodium berghei.* **B**) Parasitemia after natural transmission by mosquito bite. Note that mice bitten by mosquitoes infected with ARPC1(-) parasites are not infected. Mean +/- SD of 4 mice per group. **C**) Ookinete morphology and DNA staining. Scale bar, 5 µm. **D**) DNA intensity of ookinetes. au, arbitrary units. **E**) Mosquito infection rate as proportion of midguts carrying oocysts. **F**) Oocyst numbers per midgut at day 6 and 12 after mosquito infection. **G**) Oocyst area at day 4, 6 and 12 after mosquito infection. **H**) Oocyst morphology and DNA content at day 6 and 12 after mosquito infection. Scale bar, 10 µm. **I**) Total DNA content of oocysts 6 days after mosquito infection, normalised to the mean DNA content of Pb_473_WT. **J**) Oocyst area at day 6 after crossing Pb_473_ARPC1(-) with itself, with 47(-) (female-deficient), or with 48/45(-) (male-deficient) *P. berghei*. **K**) ARPC1-HA signal in female (top) and male (bottom) gametocytes as determined by flow cytometry. **E, F, I**) Line indicates median. **D, F, G, I, J**) Pooled data from at least 3 (J, 2) independent experiments. Grey number above x axis indicates total number of cells/midguts analysed. **C, H**) Representative images of at least 10 images taken. Statistics: **D**) Unpaired T-test, ****, p < 0.0001. **F, G, I, J**) Kruskal-Wallis Test, with Dunn’s post test. ****, p < 0.0001.

The actin-related protein 2/3 (Arp2/3) complex is a seven-subunit complex consisting of Arp2, Arp3 and five supporting subunits ARPC1-5 that together nucleate actin filaments (*8*– *10*). In the cytoplasm of most metazoa, the Arp2/3 complex mediates lamellipodia formation and endocytic trafficking, among other processes (*7*, *8*). Nuclear functions of Arp2/3 are less well studied and differ between species, but they include DNA damage repair, nucleation of spindle actin during mitosis and meiosis and chromosome capture and segregation (*18–24*). Despite the evolutionary conservation of Arp2/3 across the eukaryotic kingdom (*7*, *25*), it was assumed that the complex has been lost in apicomplexan species, except for a single subunit, annotated as ARPC1/ARC40 in *Plasmodium* (*11*, *12*). Here we discovered that *Plasmodium* ARPC1 constitutes part of a highly divergent, non-canonical Arp2/3 complex, which associates with mitotic spindles in activated male gametocytes and is essential for genome segregation into budding gametes. Disruption of the complex results in a delayed death phenotype in oocysts leading to a complete transmission block within mosquitoes.

## Results

### PbARPC1 is essential for male gamete fertility and hence mosquito stage development

Previous phylogenetic studies identified *Plasmodium* ARPC1/ARC40, from here on named ARPC1 to be consistent with the most common Arp2/3 complex subunit nomenclature, as the sole conserved subunit of the Arp2/3 complex in *Plasmodium*. In the rodent malaria parasite, *P. berghei* PbARPC1 (accession number PBANKA_0929300, PF3D7_1118800 in the human malaria parasite, *P. falciparum*) is a 41 kDa protein predicted to fold into a WD40 repeat domain with two alphahelical loops extending from the doughnut-shaped β-propeller (*26*, *27*) (**fig. S1A**). The ARPC1 protein sequence is conserved across the genus *Plasmodium* but shows less than 20% identity to other ARPC1/ARPC1 proteins from model species (**fig. S1B, C**). *Plasmodium* ARPC1 has also low sequence identity to a predicted ARPC1 homologue of *Cryptosporidium parvum*, the only other ARPC1 protein predicted to be present in apicomplexans (*11*). Tagging ARPC1 C-terminally with GFP in *P. berghei* wildtype (WT) parasites (**fig. S2A, B**) revealed that the protein is predominantly expressed in gametocytes and in ookinetes, the stage of the parasites penetrating the mosquito midgut epithelium, indicating a potential role during transmission to the mosquito (**fig. S2C**). In both stages, PbARPC1 localises to the nucleus. A weak ARPC1-GFP signal was also observed in late-stage oocysts.

To interrogate ARPC1 function we generated a knockout (KO) of PbARPC1 in the Pb_820_WT background (Pb_820_ARCP1(-), **fig. S3A, B**). Pb_820_WT expresses cytosolic RFP in female gametocytes and GFP in male gametocytes (*28*), allowing to track asexual replication and gametocyte formation by flow cytometry (**fig. S4A**). After inoculating mice with 1000 infected red blood cells (iRBC), we found no difference in asexual growth rate or formation of female or male gametes between Pb_820_WT and Pb_820_ARPC1(-) (**fig. S4B, C**). However, when testing transmission capacity of Pb_820_ARPC1(-) infected mosquitoes, we found that mice could not be infected by mosquito bites, indicating that ARPC1 is essential for parasite development in the mosquito (**Fig. 1B, Table S1**). To further investigate the specific developmental block within mosquitoes and to facilitate phenotyping *Plasmodium* across multiple life cycle stages, we created another PbARPC1 KO in the reporter line Pb_473_WT, which expresses RFP constitutively throughout the life cycle (**fig. S3A, C**). We first probed the formation of ookinetes, the motile zygote that forms in the mosquito midgut after a blood meal. Pb_473_ARPC1(-) and Pb_820_ARPC1(-) ookinetes developed normally and exhibited- WT-like morphology and motility (**Fig. 1C, fig. S4D, E**). Yet curiously, we noted that the DNA content of ookinete nuclei was reduced by about 30% in Pb_473_ARPC1(-) compared to Pb_473_WT (**Fig. 1D**). Ookinetes were fully infective, as infecting mosquitoes with Pb_473_ARPC1(-) led to WT-like infection rates (prevalence) and oocyst numbers at early development (6 days post infection (dpi) (**Fig. 1E, F)**. However, oocyst numbers dropped significantly during later oocyst development (12 dpi) (**Fig. 1F**). Notably, none of the oocysts sporulated, and we detected no sporozoites in midguts or salivary glands. In line with this observation, none of the mice bitten by Pb_473_ARPC1(-) infected mosquitos became infected (**Table S1**). We imaged WT and KO oocysts at 4, 6, and 12 dpi and found Pb_473_ARPC1(-) oocysts to be significantly smaller compared to WT at all time points (**Fig. 1G, H**). Pb_473_ARPC1(-) oocysts also exhibited a lower DNA content than WT oocysts (**Fig. 1H, I**). Complementing the Pb_473_ARPC1(-) line by reintroducing the *ARPC1* gene into the same locus (**fig. S3A, D**) fully restored oocyst size, sporulation, and parasite transmission, confirming that the observed phenotype is caused by deletion of the *ARPC1* gene (**fig. S5, Table S1**). In conclusion, our data demonstrate that PbARPC1 is required for normal oocyst growth and sporozoite development, and deletion of ARPC1 leads to a complete block in transmission.

In the mosquito, *Plasmodium* parasites undergo sexual replication, and many defects during mosquito stage development are inherited from one sex only (*14*, *29–31*). To investigate if ARPC1 function is sex-specific, we crossed Pb_473_ARPC1(-) parasites with either Pb47(-) parasites that do not produce fertile females (*31*) or with Pb48/45(-) parasites that do not produce fertile males (*30*). Only crossing with female-deficient (male-competent) Pb47(-) restored oocyst size at 6 dpi, while oocysts of the cross with male-deficient (female-competent) Pb48/45(-) parasites remained small (**Fig. 1J**). ARPC1 is thus required for male, but not female fertility. To test whether PbARPC1 expression is restricted to males, we tagged PbARPC1 with an HA-tag in the gametocyte reporter line Pb_820_WT (**fig. S6A, B**). Both RFP-positive female and GFP-positive male gametocytes expressed ARPC1-HA in the nucleus (**fig. S6C**), confirming the original observation with the PbARPC1-GFP line (**fig. S2C**). However, we detected both by IFA and by flow cytometry a stronger ARPC1-HA signal in males compared to females (**Fig. 1K, fig. S6C, D**), indicating a higher protein expression in the male lineage, in line with the male phenotype of PbARPC1(-).

### PbARPC1 is required for correct chromosome condensation and segregation into male gametes

In the mosquito midgut, male gametocytes undergo gametogenesis, during which they rapidly replicate their DNA three times by endomitosis and form eight axonemes in the cytoplasm, before finally eight flagellated gametes bud of the parent cell (**Fig. 2A**). Imaging ARPC1-GFP during male gametogenesis revealed that ARPC1 relocalises from the nucleoplasm at 0 minutes post activation (mpa) to an area surrounding the mitotic spindle at 3 mpa (**Fig. 2B**, full figure in **fig. S7**). ARPC1 then followed the spindle dynamics, localising to two spindles at 7-8 and to four spindles at 12 mpa. At 15 mpa, ARPC1 was found either at eight distinct foci surrounding the homogeneous DNA signal, or within the residual body beneath eight condensed DNA foci or the emerging flagella. To investigate if spindle formation, DNA replication or DNA condensation are impaired in ARPC1(-), we next imaged Pb_473_ARPC1(-) gametocytes at 3 mpa and at 15 mpa. At 3 mpa, both Pb_473_WT and Pb_473_ARPC1(-) gametocytes contained a spindle (**Fig. 2C**). At 15 mpa, Pb_473_ARPC1(-) gametocytes formed axonemes and replicated their DNA (**Fig. 2D**). However, we observed differences in DNA localisation. In Pb_473_WT gametocytes, about half of all male gametocytes contained an enlarged nucleus with a homogeneous DNA distribution, in about 30% of cells the DNA was condensed into eight foci, and 20% of cells were exflagellating, with DNA localising to the emerging flagella (**Fig. 2D, E**). In the remaining cells, DNA was localised in an irregular pattern, likely representing an intermediate step in the process of DNA segregation (i.e., between homogeneous distribution and the condensed localisation). In contrast, only few Pb_473_ARPC1(-) gametocytes had condensed DNA foci while the majority exhibited an irregular DNA pattern (**Fig. 2J, K**). In exflagellating Pb_473_ARPC1(-) gametocytes, a large proportion of the DNA was retained in the residual body and only a part of the DNA localised to the flagella. Altogether, our data suggests that ARPC1 is not required for spindle or gamete formation but involved in the process of DNA condensation right before budding of the flagellated gamete.

**Figure 2:**
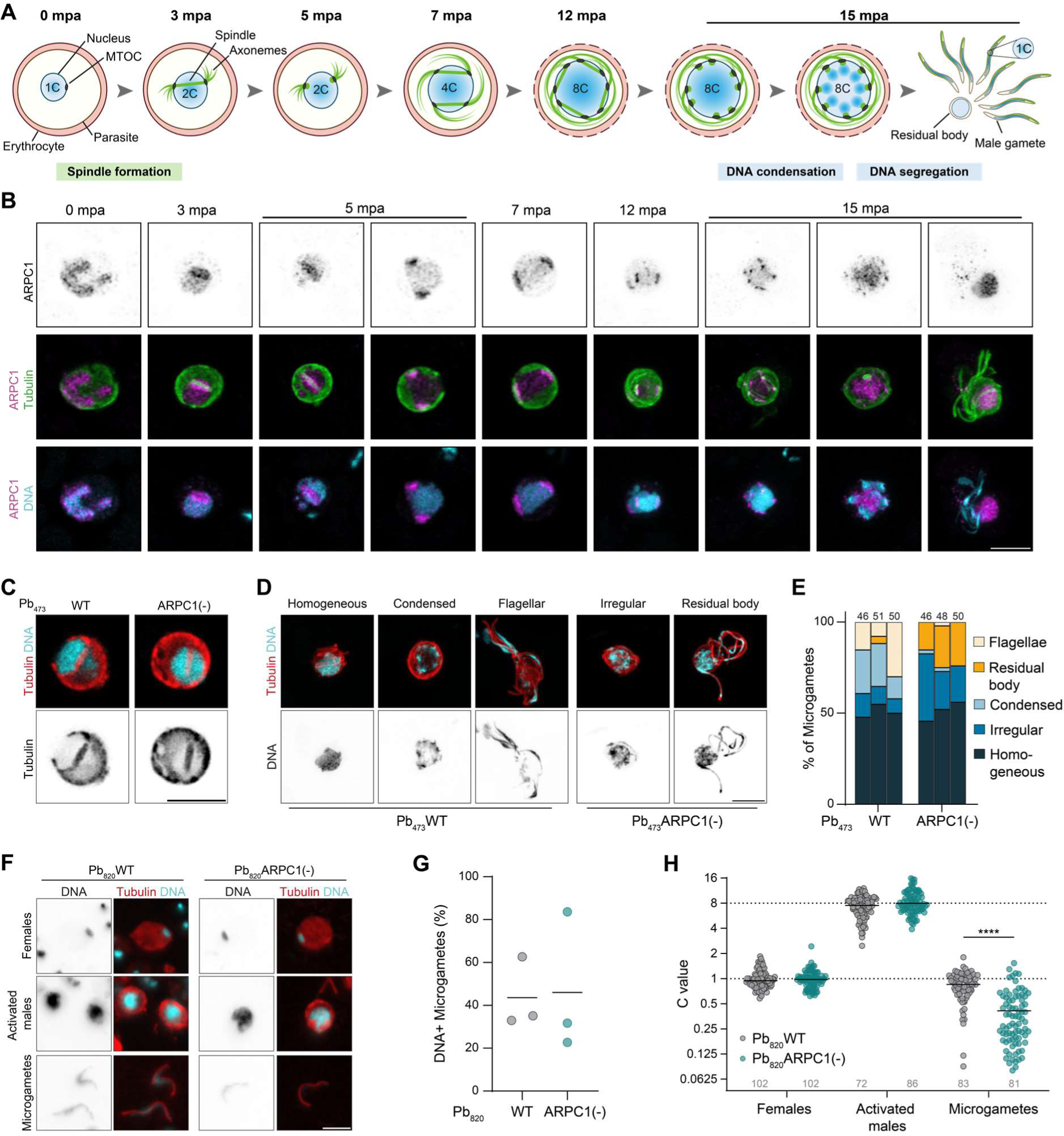
**ARPC1 mediates DNA segregation into male gametes. A**) Scheme of male gametogenesis. mpa, minutes post activation. **B**) ARPC1-GFP localisation during gametogenesis. 0-7 mpa: single slice. 12-15 mpa: Maximum Z projection. Scale bar, 5 µm. **C**) Spindle formation in activated male gametes 3 mpa. Scale bar, 5 µm. **D**) DNA localisation in activated male gametocytes 15 mpa. Scale bar, 5 µm. **E**) Quantification of DNA localisation observed in activated male gametocytes 15 mpa. Each bar represents an individual biological replicate. **F**) Images of females, activated males and microgametes 20 mpa. Scale bar, 10 µm. **G**) Proportion of microgametes with detectable DNA signal. **H**) DNA content of females, activated males and microgametes, normalised to the mean DNA content of females imaged on the same slide. Dashed line, expected C value of activated males (8C) and microgametes (1C). Pooled data from three independent experiments. Grey number above x axis indicates total number of cells analysed. **B-D, F**) Representative images from at least 5-10 image series. Statistics: **H**) Kruskal-Wallis test, Dunn’s post test. ****, p<0.000 Pb_473_ARPC1(-), with one microtubule even extending into the nuclear envelope, the reconstruction demonstrated that basal body duplication and spindle formation has occurred in Pb_473_ARPC1(-) ookinetes despite a decreased DNA content. Thus, while ARPC1 is essential for DNA segregation during male gametogenesis, subsequent sub-haploid gametes are still fertile and ARPC1(-) parasites are able to undergo meiosis and ookinete formation.

To investigate if the change in DNA localisation also results in defects in DNA segregation into the eight progeny cells, we labelled fixed gametocytes 20 mpa for DNA and tubulin and quantified the DNA content of activated male gametocytes (strong nuclear signal and a strong tubulin signal of the axonemes wrapped around the nucleus) and free male gametes (single tubulin-positive axoneme) (**Fig. 2F-H**). DNA signal was normalised to that of female gametocytes/gametes (single nucleus and diffuse tubulin signal) on the same slide. Notably, only about 30-60% of all microgametes contained any detectable DNA signal, supporting previous notions that DNA segregation into male gametes is an error-prone process (**Fig. 2G**) (*32*, *33*). Activated male gametocytes of both Pb_820_WT and Pb_820_ARPC1(-) lines had the expected octoploid DNA content (**Fig. 2H**). DNA-positive WT microgametes had an average DNA content of 1C, indicating that DNA segregation during exflagellation is an all-or-nothing process. In contrast, while for Pb_820_ARPC1(-) microgametes, the proportion of DNA-positive microgametes did not differ from WT, their DNA content was reduced to approximately 0.4C (**Fig. 2G, H**). Deletion of ARPC1 hence results in sub-haploid microgametes that contribute less than one complete genome to the zygote, suggesting that ARPC1 is responsible for proper DNA segregation into the developing male gametes.

In the midgut, male gametes fertilise female gametes to form a zygote that within four hours undergoes DNA replication and meiosis to then develop into the ookinete (**fig. S8A**) (*15*). Accordingly, we found in a time course experiment that in Pb_820_WT, the DNA content roughly doubles from non-activated gametocytes to zygotes at 1 hours post activation (hpa), after fertilisation occurred, and then further increases to 4C at 4 hpa (after meiosis) to remain at that level in ookinetes (**fig. S8B**). In contrast, in Pb_820_ARPC1(-) parasites, the C value only increased to 1.2C after fertilisation, consistent with the decreased DNA content provided by the male gamete to the zygote. Nevertheless, the DNA content approximately doubled to about 2.8C at 4 hpa and remained at this level in ookinetes. To further investigate if nuclear architecture or meiosis is affected by the deletion of PbARPC1, we imaged mature Pb_473_WT and Pb_473_ARPC1(-) ookinetes by EM tomography and reconstructed their nuclei in 3D (**fig. S8C**). In both Pb_473_WT and Pb_473_ARPC1(-), four spindle pole bodies together with four hemispindles were present. While we noted that spindle microtubules appeared longer in

### ARPC1 is part of a functional non-canonical Arp2/3 complex

ARPC1 facilitates the assembly of the Arp2/3 complex in most eukaryotic species (*8*, *25*). However, the Arp2/3 complex is postulated to be absent in *Plasmodium* (*11*, *12*) and PbARPC1 has low similarity to canonical ARPC1 proteins (**Fig. S1B, C**). To further investigate the function of PbARPC1 and identify putative interaction partners, we performed immunoprecipitation of ARPC1-GFP from purified non-activated and activated gametocytes. PbGFP_con_ gametocytes that constitutively express nucleocytosolic GFP served as control (*34*). While in non-activated PbARPC1-GFP gametocytes, only ARPC1 was identified to be significantly enriched, we found five additional proteins to be enriched in PbARPC1-GFP activated gametocytes compared to PbGFP_con_ (**fig. S9A, Fig. 3A, Table S2**). These proteins included two actin-like proteins, annotated as Alp5a (PBANKA_0811800) and Alp5b (PBANKA_1007500), two proteins of unknown function (PBANKA_1014200 and PBANKA_1229300), and the apicomplexan-specific kinetochore protein 7 (AKiT7, PBANKA_0612300) (*35*) (**Fig. 3A**). All the identified proteins are conserved across *Plasmodium* and are specifically expressed in male gametocytes according to the malaria cell atlas, a single cell RNA-seq resource (**fig. S9B)** (*36*).

**Figure 3:**
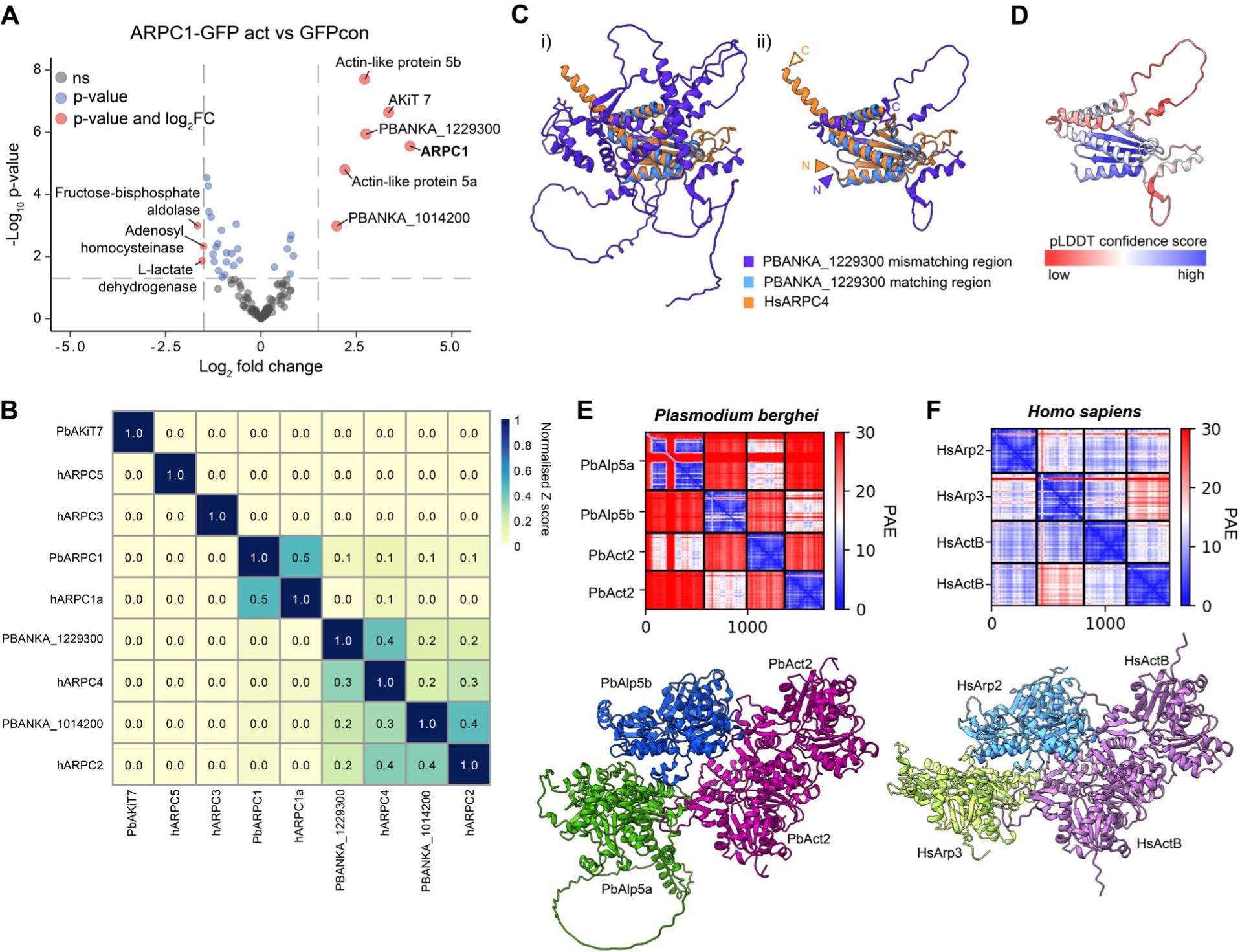
**ARPC1 constitutes part of a non-canonical Arp2/3 complex. A**) Enriched proteins after co-immunoprecipitation of ARPC1-GFP vs GFPcon from activated gametocytes followed by mass spectrometry. **B**) Structural comparisons of human ARPC subunits to *Plasmodium* ARPC candidates using DALI. Z-scores are normalized per row to the Z score of comparison to self to account for different protein sizes resulting in different maximal Z scores. **C**) Structural comparison of PBANKA_1229300 predicted structure to human ARPC4. Sub-panel i shows entire PBANKA_1229300, ii shows close-up to C-terminal region. Light blue: structural region matching to hARPC4, dark blue, structure not matching. C and N-terminus indicated in ii. **D**) PBANKA_1229300 C-terminus structural prediction colored by pLDDT confidence score indicating high confident prediction in regions aligning to hARPC4. **E**, **F**) First rank of multimer prediction of **E**) PbAlp5a, PbAlp5b and two subunits of PbAct2 and **F**) hArp2, hArp3 and two subunits of hActB. Top: Matrix showing predicted alignment error (PAE) of interaction, bottom: Predicted structure of the complex. Note the similar interaction interphase between PbAlps and PbAct2 and hArps and hActB, respectively.

Structure prediction of all identified proteins using Alphafold (*26*, *27*) suggests that Alp5a and Alp5b are structurally similar to human Arp2 and Arp3, respectively (**fig. S10A, B**). We also compared the predicted structures of ARPC1 and the two unknown *Plasmodium* proteins to those of the human ARPC subunits using the DALI algorithm (*37*). We found not only structural conservation between ARPC1 and hARPC1 as suggested by previous annotations (*11*), but also between PBANKA_1014200 and hARPC2 as well as between PBANKA_1229300 and hARPC4 (**Fig. 3B, fig. S10C-E**). Of note, the structural relationship between the *P. falciparum* homologue of PBANKA_1014200, PF3D7_1430500, and hARPC2 has been recently described, too (*2*). Intriguingly, we observed that PBANKA_1229300 contains an additional domain. While its C-terminal domain is predicted with high confidence by Alphafold and aligns very well with the structure of HsARPC4, the large N-terminal domain of around 40 kDa in size is predicted with very low confidence and consists of disordered loop regions with little secondary structure (**Fig. 3C, D**). In canonical Arp2/3 complexes, Arp2 and Arp3 nucleate the new actin filament by interaction with actin monomers (*38*). We generated Colabfold structure prediction models of Alp5a and Alp5b in complex with two subunits of the *Plasmodium* gametocyte-specific actin isoform, actin 2 (**Fig. 3E**) (*39*, *40*). Notably, this analysis predicted a direct interaction between each of the Alps with one actin subunit, and the assembled tetramer resembled closely the complex predicted for hArp2, hArp3 and two subunits of hActB (**Fig. 3E, F**). Including PbARPC1 itself, we thus identified structural homologues to five out of seven subunits of the Arp2/3 complex, including the core proteins Arp2 and Arp3, which suggests the presence of a non-canonical, minimalistic Arp2/3 complex in *Plasmodium*.

To test if the identified ARPC1 interaction partners form a complex and function in a similar manner as PbARPC1, we first tagged PbARPC2 (PBANKA_1014200) internally with YFP (**fig. S11A, B**). Imaging PbARPC2-YFPint revealed a gametocyte-specific expression and a localisation mirroring that of ARPC1: After activation, PbARPC2 relocalises from the nucleoplasm to the spindle, to ultimately retreat to the residual body upon DNA condensation (**Fig. 4A**). We also generated a knockout line of Alp5b (PbArp3) in Pb_473_WT (**fig. S11C-D**). As for ARPC1(-) parasites, this parasite line showed no decrease of blood stage growth or gametocyte production but imaging of activated Pb_473_Alp5b(-) gametocytes and gametes revealed a significant decrease in the DNA content of male gametes (**Fig. 4B, fig. S11E**).

**Figure 4:**
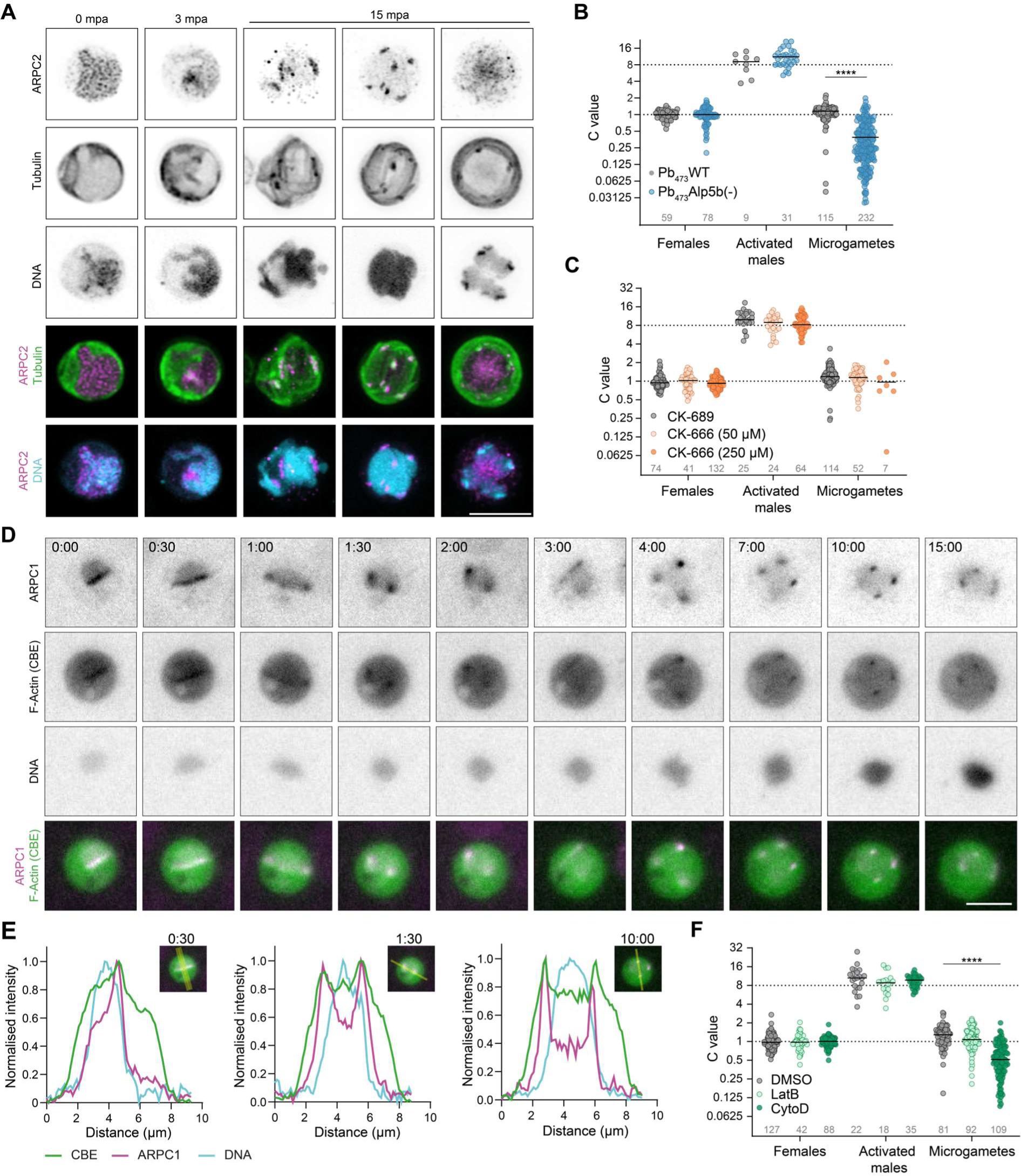
**Arp2/3 subunits and actin colocalize to the spindle and are essential for male DNA segregation. A**) Localization of ARPC2-YFP in non-activated and activated gametocytes. **B**) DNA content of females, activated males, and microgametes of Pb_473_Alp5b(-). **C**) DNA content of females, activated males, and microgametes in presence of 250 µM CK-689 (inactive control), 250 µM CK-666 or 50 µM CK-666. **D**) Live-cell imaging of activated PbARPC1-mSc/CBE male gametocyte. Time points in minutes after start of the movie indicated in upper left corner. Scale bar, 5 µm **E.)** Intensity profile plots of cross sections of selected images of the time series shown in D. Intensity was normalized to minimal and maximal value per channel. Insets show images and line used for generating the profile plots. **F**) DNA content of females, activated males, and microgametes in presence of DMSO (solvent control), 1 µM Latrunculin B (LatB) or 1 µM Cytochalasin D (CytoD). **B, C, F**) Values normalised to the mean DNA content of females imaged on the same slide. Dashed line, expected C value of activated males (8) and microgametes (1). Pooled data from at least 3 independent experiments. Grey numbers above x axis indicate total numbers of cells analysed. Statistics: Kruskal-Wallis test, Dunn’s post test. ****, p<0.0001.

These findings thus demonstrate that ARPC1 interaction partners phenocopy the function of PbARPC1 and suggest that they indeed interact in an Arp2/3-like complex to mediate male DNA segregation. We next treated gametocytes with CK-666, a drug known to target canonical Arp2/3, or its non-active control CK-689 (**Fig. 4C, fig. S11F**). Treating gametocytes during activation with 50 µM CK-666 did not affect exflagellation or the proportion of microgametes containing DNA. At a higher concentration of 250 µM, however CK-666 drastically impaired overall exflagellation rates, and the few gametes formed often did not contain DNA (**fig. S11F**). However, DNA-positive microgametes were haploid, indicating that CK-666 did not affect DNA segregation in the same manner as deletion of Arp2/3 components does (**Fig. 4C**). These data suggest that the *Plasmodium* Arp2/3 complex is highly diverse and thus not bound by the canonical Arp2/3 inhibitor.

Canonical Arp2/3 nucleates actin (*8*, *9*), and structural modelling predicted that Alp5a (PbArp2) and Alp5b (PbArp3) can interact with *Plasmodium* actin (**Fig. 3E**). To visualize actin filaments, we expressed an actin-targeting chromobody fused to GFP-emerald (CBE) (*41*, *42*) together with mScarlet-tagged ARPC1 in *P. berghei* (**fig. S12A, B**). Strikingly, live imaging showed dynamic co-localisation of ARPC1 and F-Actin throughout the entire three rounds of endomitosis (**Fig. 4D, E fig. S12C)**. Colocalisation of ARPC1 and F-Actin persisted for all three rounds of endomitosis. To investigate if actin dynamics are important for DNA segregation, we treated gametocytes in presence of cytochalasin D, an inhibitor of actin polymerisation that also impacts motility of *Plasmodium* ookinetes and sporozoites, or latrunculin B, reported to not interfere with *Plasmodium* actin dynamics (*42*). Strikingly, cytochalasin D treatment led to the formation of sub-haploid gametes, while DNA replication in male gametocytes and overall proportion of DNA-positive gametes were not affected (**Fig. 4F, fig. S12D**). In contrast, latrunculin B treatment did not affect exflagellation and DNA segregation. Taken together, the co-localisation of ARPC1 with F-actin and the sensitivity of male DNA segregation to cytochalasin D suggest that the divergent *Plasmodium* Arp2/3 complex indeed nucleates nuclear F-actin to facilitate genome segregation.

## Discussion

Early mosquito stages are a major bottleneck in the *Plasmodium* life cycle and thus a prime target for novel transmission-blocking intervention strategies (*43*). They also provide a large scope for discovery of divergent eukaryotic biology (*14*). Here we have identified a non-canonical *Plasmodium* Arp2/3 complex that is essential for DNA segregation during male gametogenesis in the mosquito (**Fig. 5**). The Arp2/3 complex localises to the spindle during male endomitosis and remains in the residual body upon gamete formation and exflagellation (**Fig. 5A**). In absence of the Arp2/3 complex subunit ARPC1, male gametes are sub-haploid, and while they can still fertilise females to form motile ookinetes, ARPC1(-) parasites arrest in a delayed-death like manner at the oocyst stage, leading to a complete block in transmission. Similarly, deletion of Alp5b results in formation of sub-haploid gametes. ARPC1 colocalises with F-actin and inhibiting actin polymerisation phenocopies ARPC1 deficiency. We therefore conclude that *Plasmodium* Arp2/3 nucleates actin, much like canonical Arp2/3 complexes. *Plasmodium* encodes for two actin isoforms, actin 1 and actin 2. Actin 2 is essential for male gametogenesis, localises both to the cytoplasm and the nuclear spindle and in its absence, gametocytes do not exflagellate (*39*). While complementation of actin 2 with actin 1 restores gamete formation and exflagellation, parasites still arrest in early oocysts, strongly suggesting that actin 1 complements cytoplasmic, but not nuclear functions of actin 2 (*44*). It is thus likely that Arp2/3 nucleates polymerisation of actin 2 along the spindle during male gametogenesis. Like Arp2/3, Actin 2 is only found in *Plasmodium* and not in other apicomplexans suggesting a specialised function of this actin isoform with the Plasmodium Arp2/3 complex.

**Figure 5:**
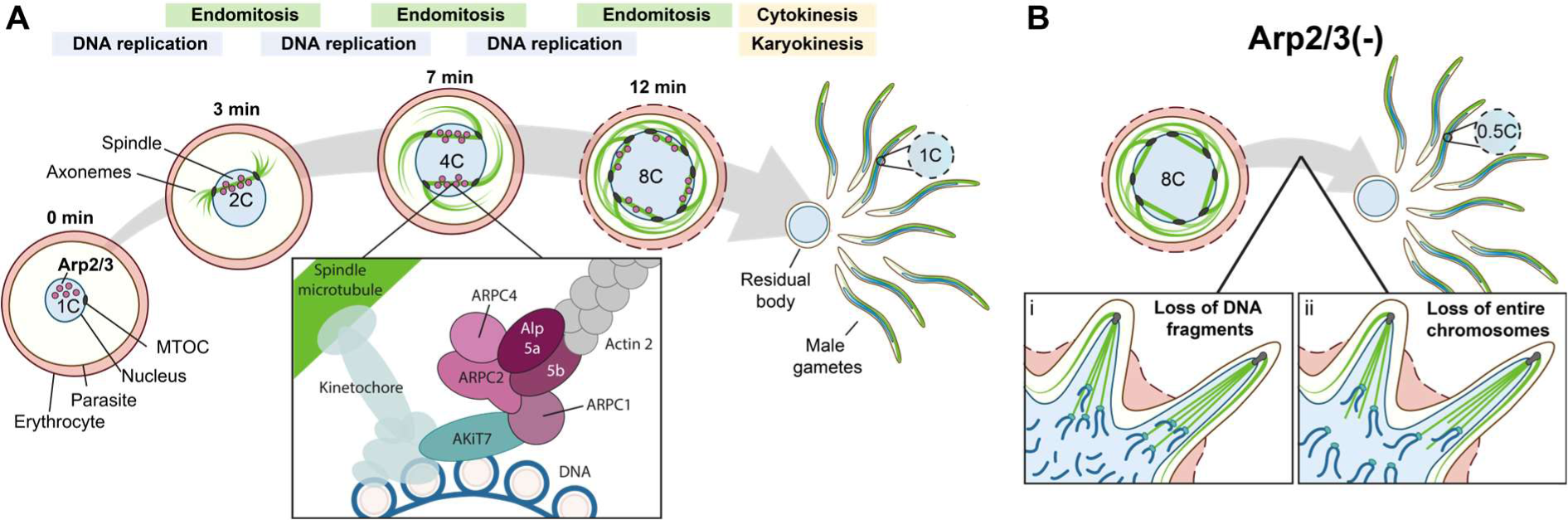
Model of *Plasmodium* Arp2/3 complex function. A) Gametogenesis in wildtype parasites. Possible localization and interactions of non-canonical Arp2/3 complex indicated. Note that directionality of actin filaments is unclear. **B**) Models for two possible causes of DNA segregation errors in absence of a functional Arp2/3 complex.

Without Alp5b or ARPC1, parasites form male gametes containing only half of the genome, i.e., the equivalent of approximately seven instead of 14 chromosomes. Two mutually compatible mechanisms could underlie *Plasmodium* Arp2/3 function (**Fig. 5B**). Canonical Arp2/3 is important for DNA damage repair and to safeguard replication forks (*18*, *19*, *45*). *Plasmodium* Arp2/3 may perform a similar function during male gametogenesis, as the three rapid rounds of DNA replication are possibly highly prone to replicative stress and DNA breaks. *Plasmodium* Arp2/3 could thus ensure that double strand breaks occurring during male DNA replication are repaired rapidly. In absence of Arp2/3, DNA double strand breaks might accumulate, and upon emergence of gametes, chromosome fragments might remain in the residual body. A second possibility is that Arp2/3 ensures correct attachment of kinetochores to the spindle. In absence of Arp2/3, some chromosomes might not connect to the spindle and thus remain in the residual body. The localisation of Arp2/3 components to the spindle and the interaction with the kinetochore protein AKiT7 argue in favour of this hypothesis, suggesting a unique function for the *Plasmodium* Arp2/3 complex. Intriguingly, AKiT7 has weak homology to mitotic arrest deficiency 1 (MAD1), a core component of the spindle assembly checkpoint in many organisms (*35*, *46*). Other putative spindle assembly checkpoint proteins have not been identified in *Plasmodium*, and a canonical spindle assembly checkpoint is thought to be absent (*5*, *6*, *17*). It is thus tempting to speculate that *Plasmodium* Arp2/3 together with AKiT7 constitutes a spindle-assembly-checkpoint-like mechanism that ensures that kinetochores are attached to the spindle before mitosis proceeds, yet without stalling progression of gamete formation in its absence.

A core protein connecting the kinetochores to the spindle during male gametogenesis is end-binding protein 1 (EB1) (*32*, *47*, *48*). Deletion of EB1 in *P. berghei* resembles the phenotype of ARPC1(-), with an arrest in a delayed-death like manner in early oocysts (*48*). In contrast, deletion of EB1 in *P. falciparum* and *P. yoelli* leads to the formation of gametes that were entirely deficient of DNA and incapable of fertilising females, highlighting interesting species-specific differences in DNA segregation in male gametocytes (*32*, *47*). A paternal phenotype followed by an early oocyst arrest has also been described for other *Plasmodium* parasite lines lacking proteins such as the formin-like protein MISFIT(*49*), the microtubule motor kinesin-8X (*50*), the aurora kinase Ark2 (*48*), and for a parasite line that expresses actin 1 in place of actin 2 (*44*). While their precise connection to the Arp2/3 complex or Arp2/3- mediated actin polymerization remains to be determined, a unifying feature of these paternal defects is that they do not affect ookinete development but arrest parasites only in the oocyst stage. Interestingly, immunoprecipitation of Ark2 and EB1 revealed many interacting proteins including Akit7 but not a single subunit of the Arp2/3 complex (*48*), suggesting a dynamic interplay between the kinetochores and the Arp2/3 complex.

The delayed-death-like developmental arrest of ARPC1(-) parasites in early oocysts remains puzzling. ARPC1(-) oocysts are still functionally diploid, meaning all genes are present and provided by the female genome, regardless of the nature of the missing male DNA. It is possible that the aneuploidy of the oocyst leads to activation of a yet unknown post-meiotic replication checkpoint, leading to an arrest in the cell cycle. Alternatively, gene expression levels could be affected in absence of the male genome, causing gene dosage mediated developmental defects. A set of paternally imprinted genes could also be expressed from the paternal genome only, possibly regulated through unknown epigenetic effects. Finally, the Arp2/3 complex could have a separate function during oocyst development. Yet, Arp2/3 expression was not detected in early oocysts at the time point of arrest, but only during sporogony in late oocysts. Notably, we detected ARPC1 expression in the nucleus of females and ookinetes. While crossing with a female-deficient line rescued the oocyst growth phenotype, it cannot be excluded that female-derived ARPC1 expression is essential at a later stage of oocyst development. The observed heterogeneous arrest of ARPC1(-) oocysts might also be a combination of all three possible effects.

Phylogenetic analyses across Apicomplexan parasites for Arp2/3 components and the apparent lack of branched actin in *Plasmodium* led to the prevailing assumption that the Arp2/3 complex had been lost (*11*, *12*). Here we report discovery of a non-canonical Arp2/3 complex that differs substantially at sequence, structural and functional level from canonical Arp2/3. While the canonical Arp2/3 complex consists of seven subunits, we have only identified five orthologues. These subunits differ markedly on a sequence level from their canonical counterparts, yet their structural core is conserved. Canonical Arp2/3 complexes are recruited and activated by nucleation promoting factors (NPFs) that either belong to the Wiscott-Aldrich Syndrome Protein (WASP) or the WISH/DIP1/SPIN90 (WDS) family (*7*). However, we could not identify any orthologues to these proteins in the *Plasmodium* genomes (*51*). The mode of Arp2/3 activation thus remains to be determined. Intriguingly, *Plasmodium* ARPC4 contains a long, N-terminal extension that does not align to any Arp2/3 subunit. Possibly, this extension either compensates for one of the missing subunits of Arp2/3, or it connects the complex to the spindle, compensating for missing NPFs. The discovery of *Plasmodium* Arp2/3 raises the question if non-canonical Arp2/3 complexes are present in other apicomplexan lineages. While we did not find any orthologues of Arp2/3 subunits using the *Plasmodium* orthologues as query, an ARPC1 orthologue has been annotated in *Cryptosporidium* (*11*) and we found ARPC2 orthologues by structure search (*52*) in *Toxoplasma*, *Eimeria* and *Neospora.* It is therefore likely that the Arp2/3 complex has assumed a specialised function in Apicomplexa that resulted in major sequence divergence while structural features remained conserved.

In conclusion, we present the discovery and characterisation of a non-canonical, divergent *Plasmodium* Arp2/3 complex that is essential for malaria parasite transmission to the mosquito. This study not only sheds light on the mechanisms of the unconventional cell division of male gametes, but also provides a new target for transmission-blocking interventions. While further work is required to elucidate the precise mode of action and evolution of this non-canonical Arp2/3 complex, interfering with Arp2/3 function could provide a building block to break the vicious cycle of *Plasmodium* transmission.

## Materials and Methods

### Ethics statement and mice

All experiments were performed in accordance with GV-SOLAS and FELASA guidelines and have been approved by the German authorities (Regierungspräsidium Karlsruhe) or according to the guidelines defined by the Home Office and UK Animals (Scientific Procedures) Act 1986 and approved by the UK Home Office (project license P6CA91811) and the University of Glasgow animal welfare and ethical review body. Female Theiler Original (TO) mice weighing 25-30 g were purchased from Envigo and were aged five to eight weeks at the time point of infection. Female Swiss mice weighing 20-25 g were purchased from JANVIER and were aged five to eight weeks at the time point of infection. Female C57Bl/6 mice weighing 18- 20 g were purchased from Charles River laboratories. For each experiment, mice were age-matched and were allocated randomly to each group. Mice were kept in groups of 2 to 4 mice per cage under specified pathogen-free (SPF) conditions within the animal facilities at Heidelberg University or University of Glasgow on a 12 hour light/dark cycle at 22 °C (± 2 °C) with *ad libitum* access to food and water.

### General maintenance of parasites and mosquitoes

The following *P. berghei* lines were used in this study as parental lines to generate transgenic parasites: PbWT, *P. berghei* ANKA; Pb_473_WT, a *P. berghei* ANKA line expressing RFP from the *hsp70* promoter (generous gift from K. Huges and A. Waters, University of Glasgow, unpublished); and Pb_820_WT, a *P. berghei* ANKA line expressing GFP from a male promoter and RFP from a female reporter (generous gift from K. Huges and A. Waters, University of Glasgow (*28*)). *P. berghei* parasite infections were initiated by intraperitoneal injection of cryostabilates (100 µl parasitic blood and 200 µl freezing solution [Alsever solution + 10% glycerol]). Parasitemia was monitored by Giemsa-stained blood smears. Mice were generally sacrificed by cardiac puncture after isofluorane- or ketamine/xylazine-induced full anaesthesia. Mosquitoes (*Anopheles stephensi*) were reared and maintained according to standard procedures.

### Generation of transgenic parasites

#### Cloning and preparation of plasmids for transfection

Cloning was performed according to standard procedures using enzymes purchased from NEB, unless stated otherwise. Gibson assembly was performed using either the Infusion kit (Takara Bio) or the NEB HiFi Assembly kit (NEB). All primers are listed in table S3. All final plasmids were checked by sequencing before transfection.

To tag ARPC1 with GFP by single crossover, a targeting vector was cloned by amplifying the 3’ region of ARPC1 using primers P1/P2 and cloning it via Gibson assembly into the EcoRI/BamHI-digested plasmid PbG-031 (generous gift from A. P. Waters, University of Glasgow). The final plasmid was linearized using NheI for transfection into Pb ANKA wildtype.

To tag ARPC1 by double crossover with a triple HA-tag, the cloning strategy involved three steps. First, the GFP cassette was removed from the vector pBAT-SIL6 (*53*) by digesting with PvuII/EcoRI and relegation of the backbone to obtain the plasmid pBAT-SIL6-mCherry. Then, 5’ and 3’ homology regions of the ARPC1 locus were PCR-amplified with primers P3/P4 or P5/P6, respectively and cloned via Gibson assembly into the pBAT-SIL6-mCherry vector using the restriction sites SpeI/XbaI and AvrII/HindIII, respectively to generate pBAT-SIL6-ARPC1- mCherry. Finally, this vector was digested with PmlI/HpaI and a 3xHA sequence, generated by overlap extension PCR using the primers P7/P8/P9/P10, was inserted using Gibson assembly. The final plasmid pBAT-SIL6-ARPC1-HAtag was linearised using KpnI/XbaI and transfected into Pb_820_WT.

To tag ARPC2 internally with a YFP, pBAT-SIL6-mCherry was digested with PmlI/NaeI. The 3’region of ARPC2 was amplified from genomic DNA in two fragments using primers P13/14 and P15/16, and a YFP gene was amplified from the plasmid pSYFP2-C1 (generous gift from Dorus Gadella (Addgene plasmid # 22878) using primers P17/18. Fragments were assembled via Gibson assembly to generate pBAT-SIL6-ARPC2-YFPint. The final plasmid was linearised using EcoRI to transfect into Pb_473_WT.

To generate a double-positive line which expresses mScarlet-tagged ARPC1 and the chromobody-emerald under the actin 1-promoter, we first amplified mScarlet using the primers P19/20 and cloned it into the HpaI/PmeI-digested pBAT-SIL6-ARPC1-mCherry. The resulting vector was then cut with SacI and ligated with the chromobody-emerald expression cassette, which was amplified using primers P21/22 from a previously published vector (*42*). The vector was linearised with XbaI/HindIII prior to transfection.

To delete ARPC1, the targeting vector PbGEM-290656 was obtained from the PlasmoGEM resource(*54*). The vector was linearised using NdeI for transfection into Pb_820_WT or Pb_473_WT.

To complement the PbARPC1(-) line, the ARPC1 gene was initially amplified from genomic DNA using primers P11/P12 and cloned into the BamHI/XbaI-digested vector pBAT-SIL6 to obtain pBAT-SIL6-ARPC1. In order to perform the complementation with a HA-tagged ARPC1, the ARPC1 gene was then excised again by SacII/XhoI digest and cloned into the equally digested pBAT-SIL6-ARPC1-HAtag plasmid to obtain the plasmid pBAT-SIL6-ARPC1- HAcomp. For transfection, the vector was digested with SacII/KpnI.

To delete Alp5b, 5’ and 3’ homology regions were amplified from genomic DNA using primers P23/24 and P25/26 respectively. The plasmid pBAT-SIL6-mCherry was digested with NaeI/SacI and AvrII/AatII to insert the 3’ and 5’ homology region via four-fragment Gibson assembly, resulting in the plasmid pBAT-Alp5b-KO. The final plasmid was digested using NotI for transfection into Pb_473_WT.

### Transfection

Transgenic *P. berghei* parasite lines were generated largely as previously described(*55*, *56*). For all transfections, 10 µg DNA were digested overnight, precipitated using ethanol and resuspended in 10 µl PBS. Schizonts were obtained by culturing 500 µl blood containing > 1% parasitemia in schizont medium (RPMI-1640 (REF 52400-025, gibco) supplemented with 20% FCS (REF 26140-079, gibco) and 1 µg/ml gentamycin (PAA) for about 20 h at 37 C, and were purified over a 55% Nycodenz (Axis-shield diagnostics) gradient. DNA and schizonts were mixed with 100 µl Nucleofector solution (either from the parasite transfection kit or from the human T cell Nucleofector Kit), electroporated using the Amaxa Nucleofector II device (Lonza, Köln, Germany), and immediately injected intravenously into a mouse. Transgenic parasites were selected by administering pyrimethamine (7 µg/ml, Sigma-Aldrich, Munich, Germany) to the drinking water one day after transfection. Blood-stage positive mice were bled by cardiac puncture and parasites were genotyped as described below. If correct transgenesis was observed, single clones were obtained by limiting dilution for all lines except for PbARPC2- YFP and PbARPC1-mSc/CBE. For this purpose, blood from an infected donor mouse was collected and serially diluted to contain a single parasite per 100 µl, which was injected intravenously into 4 to 6 mice. Mice were followed up for up to 14 days and bled upon reaching parasitemia over 1%. Blood was collected for genotyping and preparing cryostabilates. One to two clones of each transgenic line were used for further phenotypic analysis.

To obtain a marker-free line by negative selection, a Pb_473_ARPC1(-)-infected mouse was treated with 5-fluorocytosine in the drinking water (1 mg/L, Sigma Aldrich). Upon reappearing of parasites in the blood, loss of the human dihydrofolate reductase (hDHFR) selection cassette was verified by genotyping as described below and a single clone was obtained by limiting dilution.

### Genotyping

For genotyping of both *P. berghei*, blood containing at least 1% parasitemia was lysed using 0.093% saponin and the parasite pellet was resuspended in 200 µl PBS. Genomic DNA was isolated using the DNeasy Blood & Tissue Kit (Qiagen) according to manufacturer’s instructions. Parasites were genotyped by PCR for amplification of the wildtype locus (locus) or whole locus (WL), integration (int), and negative selection (ns) sites using the following primer sets: PbARPC1-GFP: P27/28 (locus), P27/29 (int); PbARPC1-HA and PbARPC1- mSc/CB-EME: P27/28 (locus), P27/30 (int); PbARPC1(-): P31/32 (locus), P33/34 (int); PbARPC1(-)ns and PbARPC1(-)compl: P33/35 (WL), P33/34 (int), P36/35 (ns), P31/32 (locus). PbARPC2-YFP: P37/38 (locus), P39/40 (int), PbAlp5b(-): P41/42 (WL), P41/43 (5’int), Pb42/44 (3’int).

### Imaging localisation of Arp2/3 subunits and interaction partners

#### Blood stage, gametocyte and ookinete immunofluorescence assays

To detect ARPC1 expression in PbARPC1-GFP or in Pb_820_ARPC1-HA, we performed immunofluorescence of blood stages and ookinetes largely as described before (*57*). Mixed blood stages were obtained from highly parasitemic mice infected 3 days prior bleeding with 25*10^6^ iRBC i.p.. Schizonts were obtained by culturing 500 µl blood containing > 1% parasitemia in schizont medium (see above) for about 20 h at 37 °C. Ookinetes were obtained by infecting a mouse with 20*10^6^ iRBC i.p. and three days later culturing about 500 µl blood in 10 ml ookinete medium (RPMI supplemented with 20% (v/v) FCS, 50 µg/ml hypoxanthine, and 100 µM xanthurenic acid, adjusted to pH 7.8 – 8.0) for 20 h at 19 °C, followed by purification over a 63% Nycodenz gradient.

For the gametocyte activation time course of PbARPC1-GFP, PbARPC2-YFP, Pb_473_WT and Pb_473_ARPC1(-), mice were infected either with 25*10^6^ iRBC i.p. and bled 3 days later, or with 15*10^6^ iRBC i.p. and bled 4 days later. The blood was immediately stored at 37 °C and 100 µl blood were transferred to 400 µl pre-warmed schizont medium for the non-activated sample. To activate gametogenesis, 100 µl blood were incubated with 400 µl ookinete medium at 19 °C. At indicated time points (0, 3, 5, 7, 12 and/or 15 mpa), gametogenesis was stopped by adding fixative.

All cell suspensions were fixed by adding equal volume of 4% PFA/0.0075% glutaraldehyde in PBS and incubating for 30 min at 37 °C. Cells were washed once in 1 ml PBS and permeabilised for 15 min at RT in 125 mM glycine/0.1% Triton-X-100 in PBS. After blocking in 3% BSA/PBS for at least 1 h at RT, cells were incubated in primary antibody for 4 h at RT or overnight at 4 °C. Cells were washed 3 times for 10 min with 1% BSA/PBS and secondary antibody was added for 1-2 h incubation at RT. Cells were washed thrice for 10 min in 1% BSA/PBS, adding Hoechst-33342 at a final concentration of 10 µg/ml to the second wash step. Cells were pelleted and 2 µl cell pellet placed on a glass slide and covered with a cover slip. Cells were imaged either on a Zeiss Axiovert 200M fluorescent microscope using a 63x objective, on a Zeiss CellDiscoverer 7 fluorescent microscope using a 50x water immersion objective, or on a Zeiss LSM900 microscope equipped with the Airyscan detector using Plan-Apochromat 63x/1.4 oil immersion objective. To detect ARPC1-HA expression in male and female gametocytes of Pb_820_ARPC1-HA, the stained samples were additionally analysed by flow cytometry on a BD FACSCelesta cell analyzer.

Primary antibodies used were rabbit-anti-GFP (1:50 dilution, Invitrogen, G10362), mouse-anti-tubulin (1:1000 dilution, Sigma-Aldrich, T5168-2ML), and rat-anti-HA (1:1000 dilution, Roche, 11867423001). Secondary antibodies used were Alexa-546-anti-rabbit (Invitrogen, A11035), Alexa-488-anti-mouse (Invitrogen, A11029), Alexa-488-anti-rabbit (Invitrogen, A11008) and Alexa-647-anti-rat (BD Pharma, 51-9006589), all at a 1:1000 dilution. For all immunofluorescence assays, matching PbWT samples were stained in parallel to control for unspecific staining of the anti-GFP or the anti-HA antibody, and no major unspecific staining was detected.

### Live cell imaging of gametocytes

For life cell imaging of PbARPC1-mScarlet-CBE, a drop of blood from highly parasitemic mouse was mixed with 3 ml ookinete medium and Hoechst-33342 at a final dilution of 10 µg/ml. Cells were immediately placed on a glass slide, covered with a cover slip and imaged on Zeiss Axiovert 200M fluorescent microscope using a 63x objective.

### Oocysts

To detect ARPC1-GFP expression in PbARPC1-GFP oocysts, mosquitoes were infected as described below and midguts were dissected at 5 and 12 days post feed. Midguts were stained with Hoechst at a final concentration of 10 µg/ml in PBS for 30 min at 37 °C and imaged on a Zeiss Axiovert 200M fluorescent microscope using a 63x objective. PbWT oocysts imaged with the same settings did not display a GFP signal.

### Sporozoites

Mosquitoes were infected with PbARPC1-GFP and salivary glands dissected and crushed in RPMI medium on day 18 as described above to isolate sporozoites. Sporozoites were seeded into four wells of an 8-well ibiTreat labtek slide (ibidi) (4*10^5^ sporozoites per well) and centrifuged for 3 min at 800xg. After 10 min incubation at RT, the supernatant was removed, and wells were washed twice with RPMI medium. Cells were fixed with 250 µl 4% PFA/0.0075% glutaraldehyde for 1 h at RT, washed thrice with PBS and permeabilised for 1 h with 0.5% Triton-X-100/PBS at RT. After additional 3 washes, cells were blocked for 1 h in 3% BSA/PBS and incubated with rabbit-anti-GFP (1:50 in 3%BSA/PBS, 200 µl/well, Invitrogen, G10362) for 1 h at 37 °C. Cells were washed three times and incubated with Alexa546-anti- rabbit secondary antibody (1:1000 in 3% BSA/PBS, 200 µl/well, Invitrogen, A11035) for another hour at 37 °C. Cells were finally washed three times with PBS, adding Hoechst at a final concentration of 10 µg/ml to the second wash step and incubating for 10 min at RT. Sporozoites were imaged on a Zeiss Axiovert 200M fluorescent microscope using a 63x objective.

### Liver stages

HepG2 cells, cultured under standard conditions in DMEM supplemented with 10% fetal calf serum, 1 mM Glutamine and 1% antibiotica-antimycotica (Gibco), were seeded into 8-well slides (nunc) at a density of 2*10^4^ cells/well. Two days later, sporozoites were isolated from mosquitoes as described above, diluted to 100 sporozoites per µl in 3% BSA/RPMI and 200 µl sporozoite mixture was added per well. Cells were incubated for one hour at 37 °C to allow invasion, and then washed once with complete DMEM. Cells were maintained in 400 µl medium per well at 37 °C. Cells were fixed 48 h post infection in 200 µl ice-cold methanol for 10 min at -20 °C, washed with 1% FCS/PBS and blocked overnight in 10% FCS/PBS. Cells were stained with rabbit anti-mCherry (abcam, ab213511) diluted 1:300 and mouse anti- PbHsp70(*58*) diluted 1:300 for 2 h at 37 °C, followed by three washes with 1% FCS/PBS and secondary staining using Alexa-546-anti-rabbit (Invitrogen, A11035), Alexa-488-anti-mouse (Invitrogen, A11029) at 1:300 dilution for 1 h at 37 °C. Hoechst was added to a final concentration of 2.5 µg/ml and cells were incubated another 15 min at room temperature before washing three times with 1% FCS/PBS and mounting with 10% glycerol/PBS. Cells were imaged on a Zeiss Axiovert 200M fluorescent microscope using a 63x objective.

### Characterisation of parasite development across the life cycle

#### *P. berghei* asexual growth and gametocyte formation

To determine parasite growth and gametocyte formation, 4 TO mice per parasite line were infected intravenously with 1000 iRBC. Parasitemia was assessed from day 4 to day 10 post infection by staining a drop of blood in DRAQ5 (1:1000 diluted in PBS) for 10 min at room temperature, washing the cells once with PBS and analysing them by flow cytometry on a MACSQuant VYB flow cytometer.

### Ookinete formation and motility assays

For ookinete formation and motility assays, TO mice were pretreated with 200 µl phenylhydrazine (6 mg/ml in PBS) intraperitoneally to induce reticulocytosis. Two days later, mice were injected ip with a parasite cryostock. Upon reaching a parasitemia above 5% (around 3 to 4 days post infection), asexual parasites were killed by supplementing drinking water with sulphadiazine (30 mg/L). Two days later, mice were bled and the gametocyte-containing blood transferred to 10 ml ookinete medium. Parasites were cultured at 19 °C for 20 to 24 hours. Ookinetes were purified using a 63% Nycodenz gradient, washed once with ookinete medium, and 2 µl of the pellet transferred onto a glass slide and covered with a cover slip. Ookinetes were imaged for 15 min with a frame rate of 20 sec at a Nikon A1R inverted confocal microscope, using a 60x oil objective.

### Mosquito infections

To infect mosquitoes, two mice were infected either with 20*10^6^ iRBC i.p. or with 2*10^6^ iRBC i.v. Three days later, mice were weighed and anaesthetised by administering ketamine/xylazine solution (20 mg/ml ketamine, 0.6 mg/ml xylazine in PBS) i.p. at 5 µl/g body weight and placed onto a mosquito cage containing approximately 400-500 mosquitoes. Mosquitoes were allowed to feed for approximately 30 min, with a change of mouse position after 15 min. After feeding, mosquitoes were immediately kept at 21 °C and 80% humidity, and fed with 10% (v/v) saccharose with 0.05% PABA and 1% (v/v) NaCl.

### Oocyst quantification and size determination

Infection intensity was assessed 4, 6, and 12 days post blood meal by dissecting 10-30 midguts in PBS. Dissected midguts were placed on a glass slide and covered with a cover slip to image using a Leica AF6000 LX or a Zeiss Axiovert 200M fluorescent microscope with a 10x objective. The prevalence of infection was determined as proportion of infected midguts and the number of red-fluorescent oocysts was counted per midgut. To measure oocyst area, infected midguts were imaged either on a Nikon A1R inverted confocal microscope using a 25x objective or on a Zeiss Axiovert 200M fluorescent microscope using a 25x objective. The area of oocysts was determined in FIJI (*59*) using a self-written macro that auto-thresholded oocysts based on their fluorescence and measured the size.

### Sporozoite numbers

To determine sporozoite production, 15 to 40 mosquitoes were dissected 17 to 18 days post blood meal, and salivary glands transferred into a tube containing 100 µl PBS. Organs were disrupted mechanically using a plastic pestle to release sporozoites. Sporozoites were counted using a haemocytometer on a Zeiss Axiostar light microscope under a 40x objective with phase contrast.

### By-bite infections

Natural transmissions by mosquito bite to mice were performed 18 to 20 days post blood meal. TO or C57Bl/6 mice were anaesthetised by administering ketamine/xylazine solution (20 mg/ml ketamine, 0.6 mg/ml xylazine in PBS) i.p. at 5 µl/g body weight and placed either on a full infected mosquito cage or on cups containing 10 female infected mosquitoes. Mice were exposed to mosquito bites for 10 to 15 min and were bitten by at least 7 mosquitoes. Mice were monitored from day 3 to day 14 post infection by daily blood smears for appearance of parasites.

### Crossing of parasite lines

To determine sex-specificity of the ARPC1(-) phenotype, we crossed Pb_473_ARPC1(-) with Pb48/45(-) (male-deficient) or Pb47(-) (female-deficient) parasites (*30*, *31*). To this end, Pb_473_ARPC1(-) parasites were mixed with equal numbers of the respective sex-specific line and two TO mice were injected i.v. with 2*10^6^ parasites each. Three days post infection, mice were anaesthetised and fed to mosquitoes as described above. Oocyst size was determined 6 days post blood meal from red-fluorescent oocysts as described above.

### Ookinete EM reconstructions

Purified ookinetes were fixed over-night at 4 °C in 2% PFA/2% glutaraldehyde/0.1 M cacodylate buffer and processed as described before (*60*). Embedded ookinetes were sectioned to 200 nm-thick sections and imaged on a transmission electron microscope at 200 kV (Tecnai F20 TEM, FEI) equipped with a Eagle 4k x 4k CCD camera (FEI). Bidirectional tilt series were acquired from -60° to + 60° in 2° increments, with a magnification of 9600x (pixel size 1.118 nm) and 14500x (pixel size 0.74 nm) for the WT and ARPC1(-) ookinete nucleus, respectively. Tomograms were reconstructed, joined and segmented using IMOD. (*61*). The inner nuclear membrane, MTOCs and spindle microtubules were rendered using 3dmod.

### DNA content of parasites

#### Quantification of DNA content

For all DNA content analysis, wildtype and mutant parasites were prepared as described below and stained in parallel using the same dilution of Hoechst 33342. Images were taken focussing on the widest area of the nucleus and analysed in FIJI (*59*) using a self-written macro. Here, nuclei were segmented using an automatic thresholding function, and the DNA signal was measured as the total fluorescence intensity of the nucleus area minus the average background fluorescence signal. Where appropriate, DNA signal was normalised to wildtype DNA signal measured in parallel or to single nucleated cells imaged on the same slide.

### Ookinete

To determine the DNA content, ookinetes were produced *in vitro* as described for motility assays, but without sulphadiazine supplementation, as we found that adding sulphadiazine to the drinking water impedes with DNA replication in wildtype ookinetes. Nycodenz-purified ookinetes were stained with Hoechst (final concentration 10 µg/ml) for 10 min and imaged on a Zeiss Axiovert 200M microscope using the 63x objective.

### Oocysts

To determine DNA content of *P. berghei* oocysts, midguts were dissected from mosquitoes 6 days post blood meal. Midguts were incubated in Hoechst (3 µM in 3% BSA/PBS) for 30 min at 37 °C and washed two times in 3% BSA/PBS. Midguts were imaged on a Nikon A1R inverted confocal microscope with a piezo z-drive using the 100x objective and Z-stack images were taken. Sum z-projections were generated before image analysis.

### Microgametes

To quantify DNA content in activated males and male gametes, mice were infected with 20 *10^6^ iRBC i.p.. Mice were bled three days later and gametocytes were purified using a 49% Nycodenz gradient kept at 37 °C. Purified gametocytes were resuspended in 500 µl ookinete medium and incubated for 20 min at 19 °C. For drug treatment, ookinete medium contained indicated concentrations of respective drugs. Cells were pelleted and blood smears on glass slides were prepared from 2 µl of the pellet. Smears were fixed with ice-cold methanol for 5 min. Cells were rehydrated for one hour in 3% BSA/PBS, stained with mouse-anti-tubulin (1:500 - 1:1000 in 3% BSA/PBS, Sigma-Aldrich, T5168-2ML), followed by secondary antibody Alexa-546-anti-mouse (Invitrogen, A111030), 1:500 – 1:1000 in 3% BSA/PBS), and Hoechst at 10 µg/ml and mounting in 10% glycerol/PBS. Cells were imaged on a Zeiss CellDiscoverer 7 fluorescent microscope using a 50x objective. Microgametes were identified by their tubulin signal and scored as DNA-positive or -negative before proceeding to determining DNA content. Images were analysed in a single-blinded manner.

DNA content during fertilisation and ookinete development.

To quantify the DNA content over time during fertilisation, mice were infected with 20 *10^6^ iRBC i.p. or 2*10^6^ iRBC i.v. and bled three days later. The collected blood was immediately transferred to ookinete medium (6 wells a 100 µl blood and 2 ml ookinete medium) and incubated at 19 °C to induce gametogenesis and fertilisation. To obtain non-activated gametocytes, 100 µl blood were kept at 37 °C and fixed immediately. At 1 h, 2 h, 4 h, 8 h and 24 h post induction each, one well containing 100 µl blood was collected and fixed by adding equal volume of 4% paraformaldehyde (PFA, EM-grade)/0.0075% glutaraldehyde (EM-grade)/PBS for 10 min at room temperature, followed by one wash in PBS. After collection of all time points, cells were stained with Hoechst (final concentration 10 µg/ml) for 15 min at 37 °C and imaged on a Zeiss CellDiscoverer 7 fluorescent microscope using a 50x objective. Female gametocytes, zygotes and ookinetes were identified based on their red fluorescence in the reporter line Pb_820_. Images were analysed in a single-blinded manner.

### Immunoprecipitation and mass spectrometry

#### Immunoprecipitation of ARPC1-GFP

ARPC1-GFP and GFP-con gametocytes were purified from the blood of highly parasitemic mice as described above. Gametocytes were divided into two samples, with one being fixed immediately (non-activated sample), and the other being activated in ookinete medium for 4 min at 19 °C. Cells were pelleted, immediately fixed by resuspension in 1% (v/v) formaldehyde for 10 min and then quenched with 0.125 M glycine in PBS for 5 min. ARPC1- GFP was co-immunopurified using the GFP-Trap Agarose Kit (gtak-20) by chromotek according to the manufacturer’s instructions. Briefly explained, the lysis buffer was supplemented with 2.5 mM MgCl_2_ and 100 U of DNaseI, and both lysis and dilution buffer were supplemented with 1x concentration of Halt^TM^ Protease and Phosphatase Inhibitor Cocktail (Invitrogen). Gametocytes were pelleted, resuspended in 400 µl lysis buffer and lysed for 60 min on ice with regular vortexing. The GFP-trap beads were added to the supernatant and incubated rotating over end for 60 min. The beads were rinsed three times with wash buffer and proteins were finally released by resuspending beads in 2x Laemmli SDS-sample buffer and denaturation at 95 °C for 5 min. Samples were separated on a SDS-PAGE to be analysed for mass spectrometry.

### In-gel tryptic digestion

Upon SDS-PAGE, coomassie stained bands (3 per analyzed sample) were manually excised from the gel. The gel pieces were washed once with 60 µL of 1:1 (v/v) 50 mM triethylammonium bicarbonate buffer (TEAB; Sigma-Aldrich, Germany) and acetonitrile (ACN; Roth, Germany), pH 8.5 for 10 min and shrunk three times for 10 min each in 60 µL ACN and washed in 60 µL 50 mM TEAB, pH 8.5. Following a reduction of proteins with 10 mM DTT (Sigma-Aldrich, Germany) in 100 mM TEAB at 57 °C for 30 min and dehydration of gel pieces, proteins were alkylated with 10 mM IAA (Sigma-Aldrich, Germany) in 100 mM TEAB at 25 °C for 20 min in the dark. Prior to protein digestion, gel pieces were washed with 60 µl 100 mM TEAB and shrunk twice for 10 min in 60 µl ACN. A total of 30 µL of 8 ng/µL in 50 mM TEAB trypsin solution (sequencing grade, Thermo-Fisher, USA) was added to the dry gel pieces and incubated 4 h at 37 °C. The reaction was quenched by addition of 20 µL of 0.1% trifluoroacetic acid (TFA; Biosolve, The Netherlands). The resulting peptides were extracted once for 30 min with 30 µl 1:1 (v/v) 0.1% TFA and ACN, followed by gel dehydration with 20 µl ACN for 20 min, and wash with 30 µl of 100 mM TEAB for another 20 min. Finally, gel was shrunk twice with 20 µl of ACN for 20 min. The supernatant from each extraction step was collected, concentrated in a vacuum centrifuge and dissolved in 15 µl 0.1% TFA. Peptides from corresponding gel lanes were combined (1-3, 4-6 etc).

### LC-MS/MS analysis

Nanoflow LC-MS^2^ analysis was performed with an Ultimate 3000 liquid chromatography system coupled to an Orbitrap QE HF (Thermo Fisher). An in-house packed analytical column (75 µm x 200 mm, 1.9 µm ReprosilPur-AQ 120 C18 material (Dr. Maisch, Germany) was used. Mobile phase solutions were prepared as follows, solvent A: 0.1% formic acid / 1% acetonitrile, solvent B: 0.1% formic acid, 89.9% acetonitrile. Peptides were separated in a 60 min linear gradient started from 3% B and increased to 23% B over 50 min and to 38% B over 10 min, followed by washout with 95% B. The mass spectrometer was operated in data-dependent acquisition mode, automatically switching between MS and MS2. MS spectra (m/z 400–1600) were acquired in the Orbitrap at 60,000 (m/z 400) resolution and MS2 spectra were generated for up to 15 precursors with normalized collision energy of 27 and isolation width of 1.4 m/z.

### Database search

The MS/MS spectra were searched against the Swiss-Prot Mus musculus (UP000000589) and Plasmodium berghei (UP000074855) protein databases and a customized contaminant database (part of MaxQuant, MPI Martinsried) using Proteome Discoverer 2.5 with Sequest HT (Thermo Fisher Scientific). A fragment ion mass tolerance was set to 0.02 Da and a parent ion mass tolerance to 5 ppm. Trypsin was specified as enzyme. Carbamidomethyl was set as fixed modification of cysteine and oxidation (methionine), and deamidation (asparagine, glutamine) as variable modifications of peptides. Acetylation, methionine-loss and combination of acetylation and methionine-loss were set as variable modifications of protein terminus. Peptide quantification was done using precursor ion quantifier node with Top N Average (n = 3) method set for protein abundance calculation.

### Data analysis

Only *P. berghei* proteins detected in at least two out of the four ARPC1-GFP-activated replicates were included in further data analysis. The data set was processed in Perseus (v2.0.10) (*62*) and R (v4.2.2). Protein abundance was log2-transformed and missing values were imputed in Perseus (width 0.3, downshift 1.8). The data was normalized by Z-transformation on each column separately before calculating fold changes. P values were calculated using a Students t-test.

### *In silico* structural analysis

Protein structure and interface predictions were modelled by AF2 multimer version 3 (*63*)using ColabFold v1.5.2 (*40*). Diverse multiple sequence alignments were assembled by MMseqs2. Structure predictions were generated with 12-20 recycles. Protein structure prediction alignments between *Plasmodium* and human orthologues were generated by the “all against all” feature of the Dali server (*37*).

### Statistics

Unless stated otherwise, experiments were repeated at least three independent times, and exact sample sizes are given in the figure legends. Where possible, images were analysed in a single-blinded manner. Error bars indicate SEM unless stated otherwise. Significant differences between samples are indicated with asterisks as follows: \**P* < 0.05; \*\**P* < 0.01; \*\*\**P* < 0.001; \*\*\*\**P* < 0.0001.

### Software

Most data was plotted and analysed in GraphPad Prism (v10.0.3), except for mass spectrometry data, which was analysed using Perseus (v2.0.10) and R (v4.2.2). Flow cytometry data was analysed using FlowJo (v10.7.2). ZEN Blue 3.01 software was used for the post-2D or 3D Airyscan processing with automatically determined default Airyscan. All images were further processed in FIJI (v2.14.0). Protein structure predictions were performed using Alphafold2 and ColabFold (v1.5.2). Structures were visualised using UCSF ChimeraX (1.6.1). EM tomograms were joined and reconstructed using IMOD.

## Supporting information

Table S2 - Mass spectrometry data

## Acknowledgments

We thank M. Reinig (Heidelberg University) and A. McIlhoney (University of Glasgow) for mosquito rearing. We thank A. P. Waters and K. Huges (University of Glasgow) for generously sharing parasite lines and plasmids; M. Ganter (Heidelberg University) and R. Douglass (Gießen University) for fruitful discussion and critical comments to the manuscript; and M. Luzarowski and S. Merker from the Core Facility for Mass Spectrometry Proteomics (CFMP) at the ZMBH Heidelberg for technical support, performing mass spectrometry, and advice regarding data analysis. We further acknowledge the microscopy support from the Infectious Diseases Imaging Platform (IDIP) at the Center for Integrative Infectious Disease Research, Heidelberg. We are thankful to S. Gold and C. Funaya and the Electron Microscopy Core Facility (EMCF) Heidelberg for support in preparing and imaging EM samples. The *Plasmodium* database PlasmoDB facilitated this work.

## Author contributions

Conceptualization: FH, MM Methodology: FH, MM, FF, MS

Investigation: FH, DJ, YS, PA, AK, KH, LD, MS

Formal analysis: FH, DJ

Visualization: FH, DJ

Funding acquisition: FH, MM, FF

Supervision: FH, MM, FF, MS

Writing – original draft: FH

Writing – review & editing: FH, MM, FF

## Data and materials availability

Raw mass spectrometry proteomics data have been deposited to the ProteomeXchange Consortium via the PRIDE partner repository (*64*) with the dataset identifier PXD046181. All other data are available in the main text or the supplementary materials. Materials are available upon request.

## Supplementary Materials

Figs. S1 to S12 Tables S1 to S3

