## Supplementary Figures and Tables for "A non-canonical Arp2/3 complex is essential for *Plasmodium* DNA segregation and transmission of malaria"

**Affiliations**

**This PDF contains:**

Fig. S1-12
Table S1
Table S3

**Other supplementary materials for this manuscript include:**

Table S2

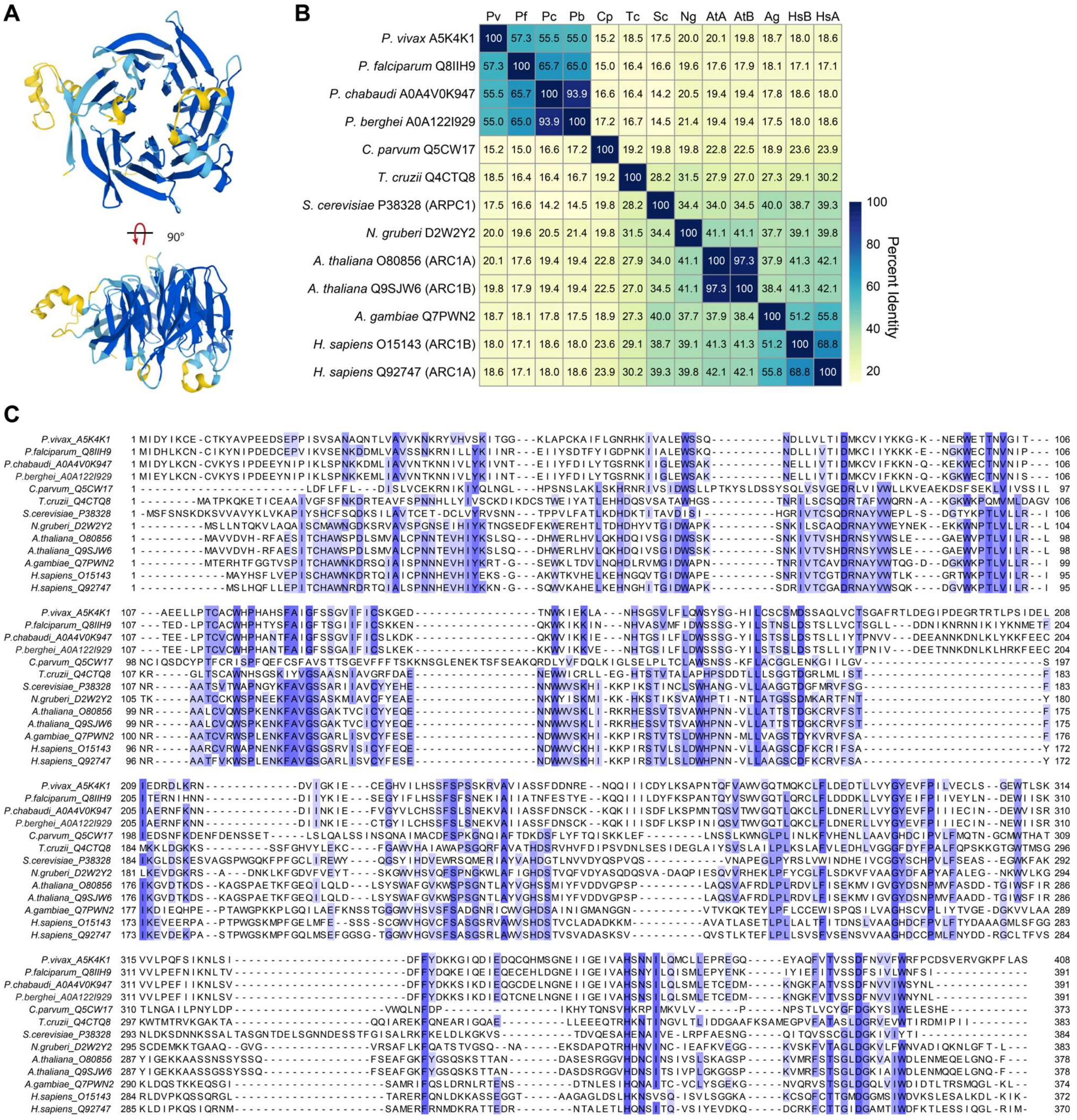


**Figure S1: ARPC1 is a *Plasmodium*-specific WD40 domain protein. A**) Alphafold prediction of *P. berghei* ARC40 structure. **B)** Percent Identity Matrix of *Plasmodium* ARC40 and the ARC40/ARPC1 subunit from other model organisms, based on Clustal Omega alignments. **C**) Clustal Omega Alignment of putative ARC40/ARPC1 sequences from *Plasmodium* and selected model organisms.

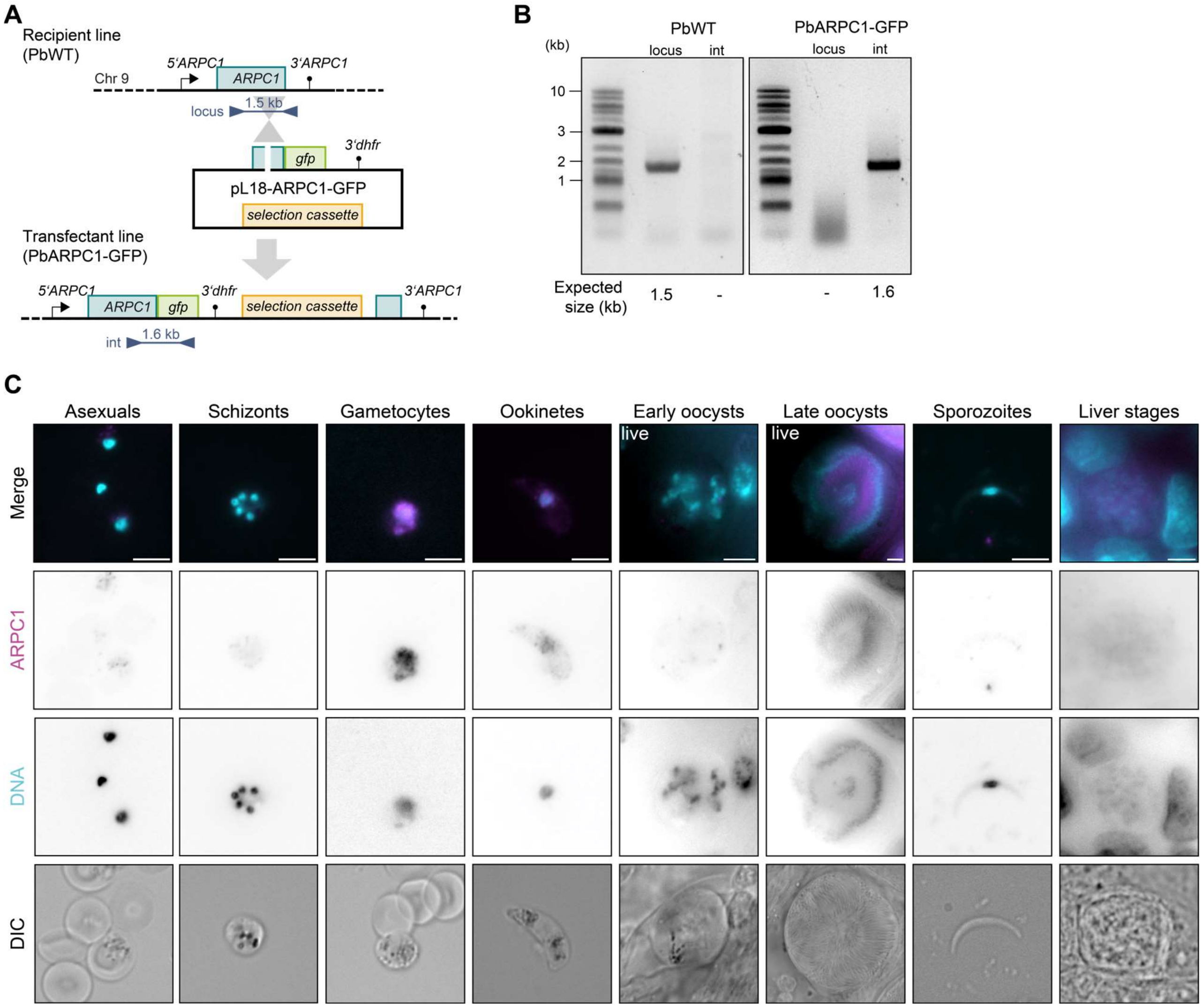


**Figure S2: Generation and localization of ARPC1-GFP**. **A**) Scheme of genetic strategy. Primers (triangles) and expected fragment sizes used for genotyping (see B) are indicated. Not drawn to scale. **B**) Genotyping PCR of PbARPC1-GFP. The expected size of the product is indicated below the gel images. **C)** Expression and localisation of ARPC1-GFP across the life cycle of *P. berghei*. Cells were fixed for immunofluorescent staining, except for oocysts, which were imaged live. Representative images of at least 10 images. Scale bar, 5 µm.

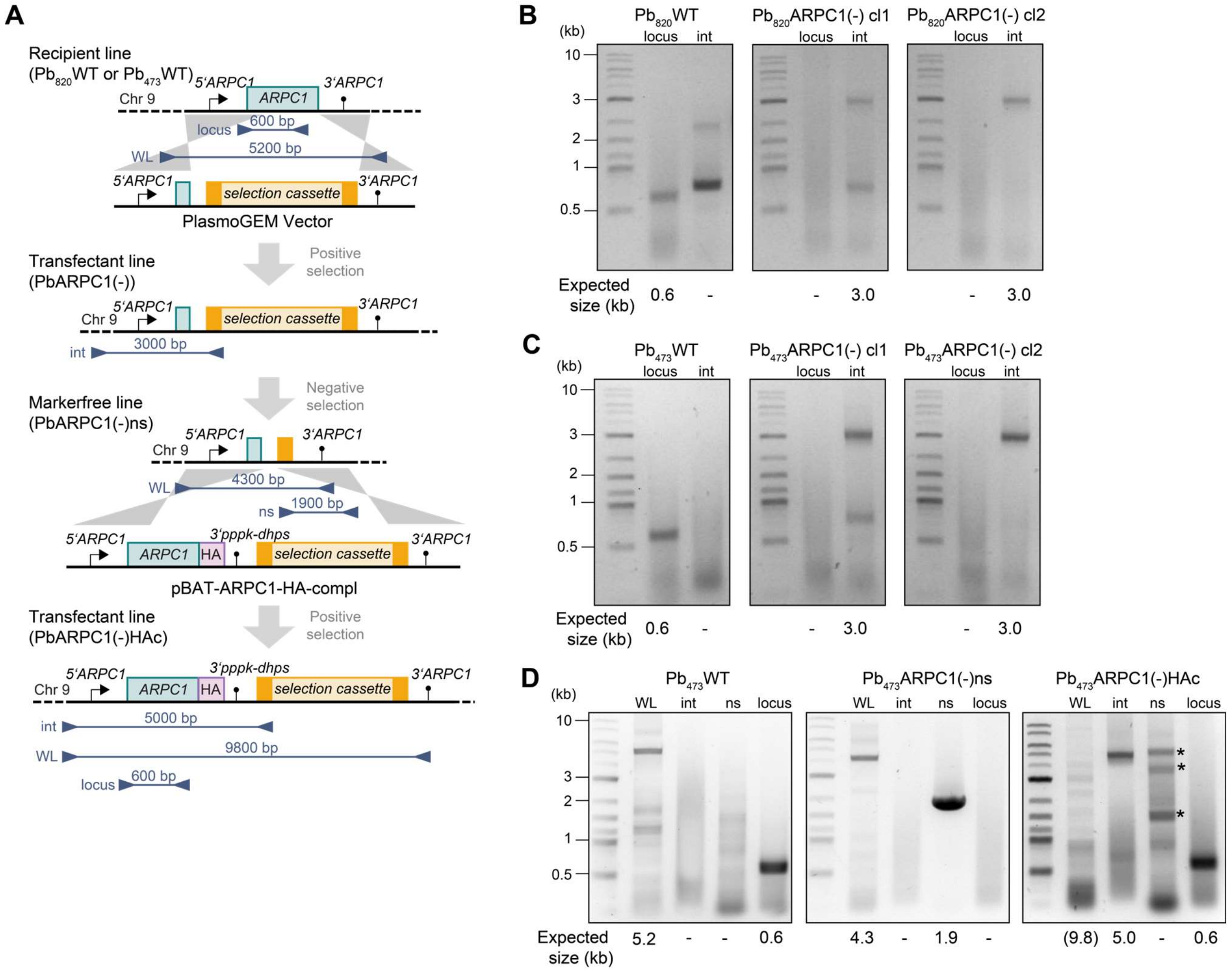


**Figure S3: Generation and characterisation of PbARPC1(-) and complementation lines**. **A**) Scheme of genetic strategy. Primers (triangles) and expected fragment sizes used for genotyping (see B-D) are indicated. Not drawn to scale. **B-D**) Genotyping PCR of **B**) $Pb_{820}$ARPC1(-), **C**) $Pb_{473}$ARPC1(-) and **D**) $Pb_{473}$ARPC1(-)HAc. Binding sites of the respective primers are indicated in A. The expected size of the product is indicated below the gel images. Note the presence of some unspecific amplification products in some PCRs (indicated with asterix).

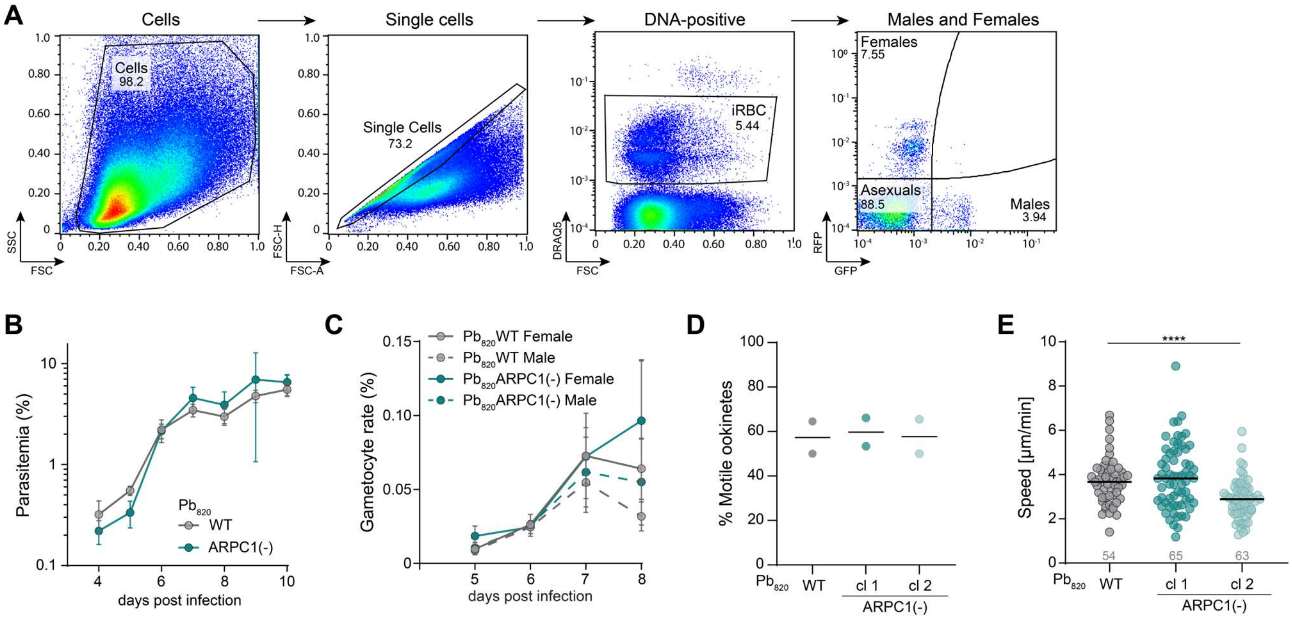


**Figure S4:** ARPC1 is not required for asexual growth, gametocyte formation or ookinete motility. **A**) Gating strategy to determine asexual growth and gametocyte formation in $Pb_{820}$ARPC1(-). **B, C**) Asexual growth (B) and gametocyte rate (C) after intravenous inoculation of mice with 1000 iRBC. Mean +/- SD of 4 mice per group. **D**) Proportion of motile ookinetes. Each data point corresponds to an independent experiment. **E**) Speed of moving ookinetes. Pooled data from two independent experiments. Grey numbers above x axis indicate total number of observed ookinetes. **E**) One-Way ANOVA, Dunn's post test. *, $p < 0.05$.

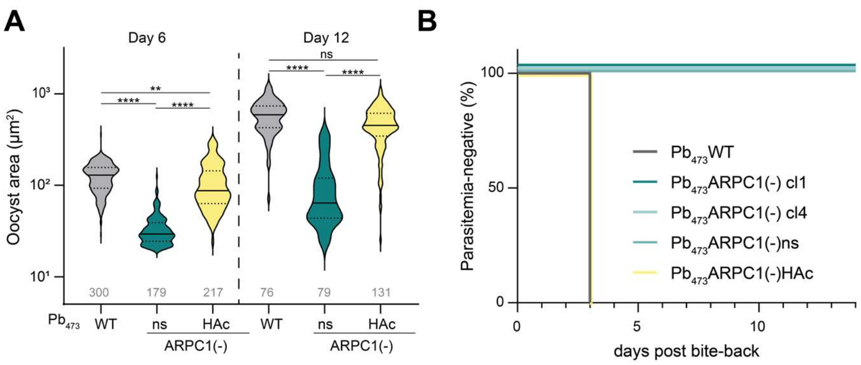


**Figure S5: Complementation of Pb$_{473}$ARPC1(-). A**) Oocyst area at day 6 and 12 after mosquito infection. Pooled data from 2 independent cages. Grey number above x axis indicates total number of cells/midguts analysed. Statistics: Kruskal-Wallis Test, with Dunn's post test. **, $p < 0.01$, ****, $p < 0.0001$. **B**) Parasite prevalence in mice after by-bite infection with 5-10 infected mosquitoes each. 3 mice per group.

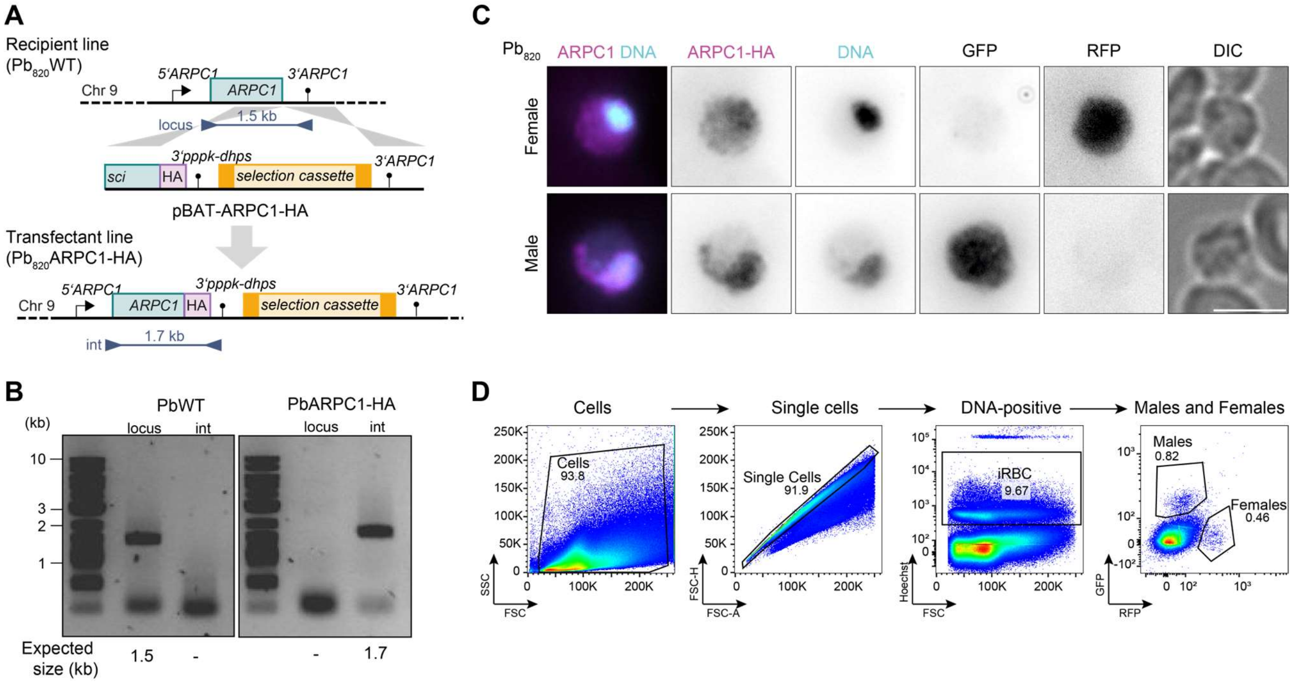


**Figure S6: Generation of Pb820ARPC1-HA**. **A**) Scheme of genetic strategy. Primers (triangles) and expected fragment sizes used for genotyping (see B-D) are indicated. Not drawn to scale. **B**) Genotyping PCR. The binding sites of the respective primers are indicated in A. **C**) ARC40-HA expression and localisation in female (RFP-positive, first row) and male (GFP-positive, second row) gametocytes. Representative widefield images of at least 10 images taken. Scale bar, 5 µm. **D**) Gating strategy to identify ARPC1-HA expression in male and female gametocytes.

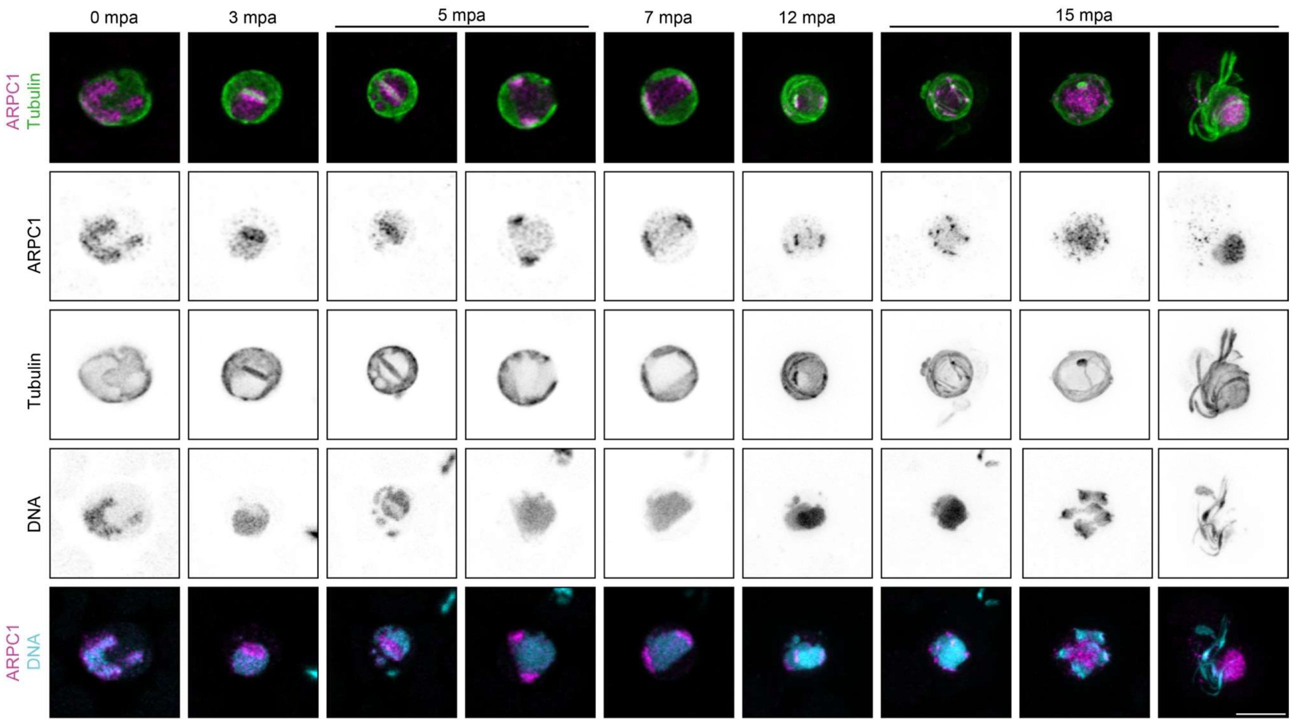


**Figure S7: Localization of ARPC1-GFP during male gametogenesis**. Activated PbARC40-GFP male gametocytes fixed at various time points after activation and stained for ARC40, tubulin and DNA (Hoechst). Shown are extended panels to Figure 3F. Representative confocal images of at least 5 images taken per time point. 0-7 mpa: Single slice. 12-15 mpa: Maximum Z projection. Scale bar, 5 µm.

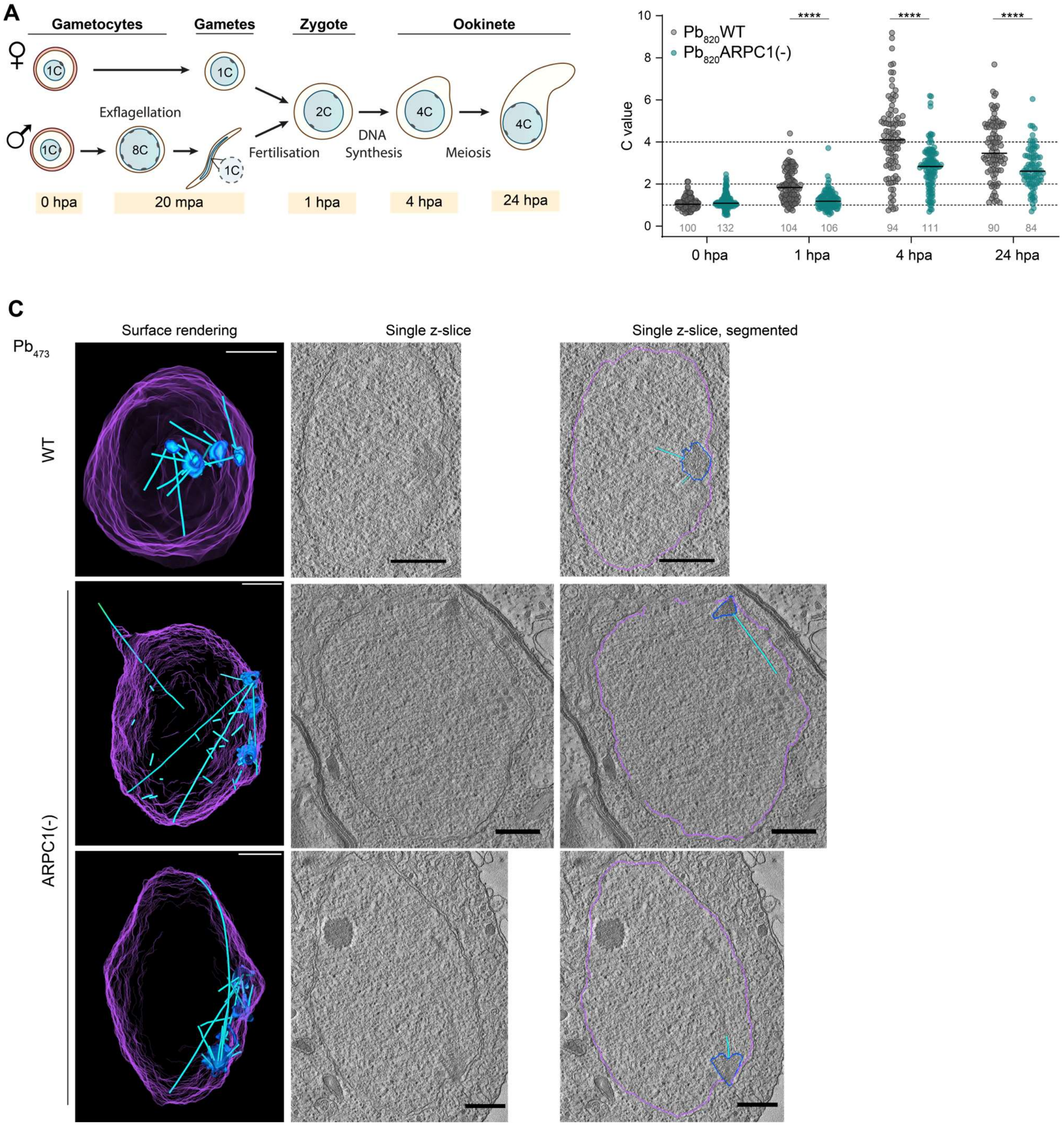


**Figure S8: Ookinete phenotype of PbARCPC1(-). A**) Scheme of *Plasmodium* sexual replication. C, C value (number of haploid genomes); mpa/hpa minutes/hours post activation. **B**) DNA content of female gametocytes, zygotes and ookinetes, normalised to the mean DNA fluorescence at 0 hpi. Dashed lines, expected C values at 0 hpi (C=1), 1 hpi (C=2) and 24 hpi (C=4). hpi, hours post induction. Statistics: Two-Way ANOVA, Šidák's post test. ****, $p < 0.0001$. **C**) 3D segmentation (first column) and example single z-slices of (second and third) of $Pb_{473}$WT and $Pb_{473}$ARC40(-) ookinete nucleus tomogram. Purple, nuclear membrane; blue, MTOC; light blue, microtubules. Scale bar 500 nm (surface rendering) and 400 nm (single sections).

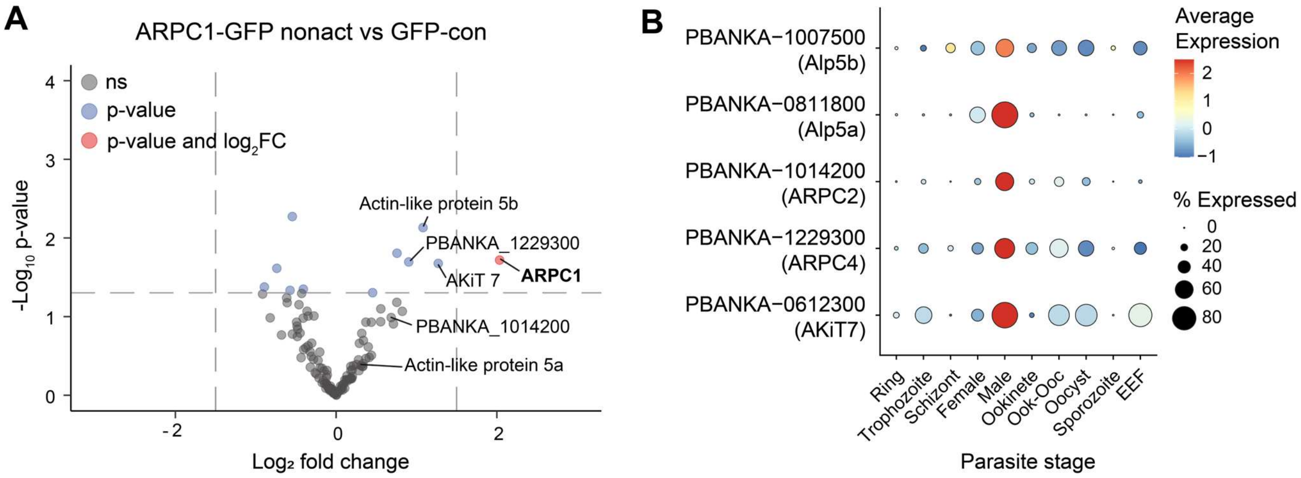


**Figure S9: Pulldown of ARPC-1 GFP. A**) Enriched proteins after pulldown of ARPC1-GFP from non-activated gametocytes. **B**) Expression of ARPC1-interaction partners across life cycle. Data reanalysed from the malaria cell atlas *(36)*.

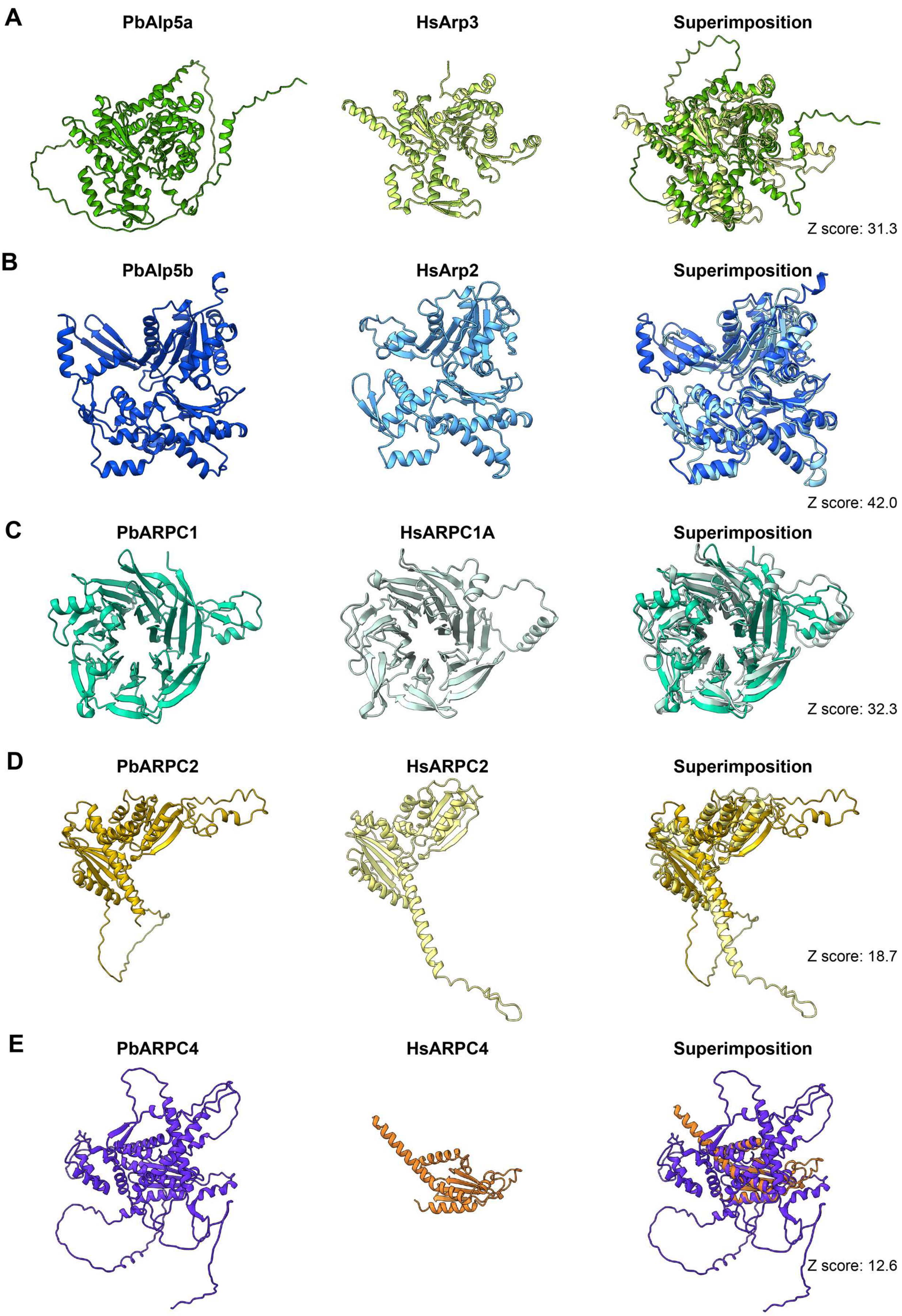


**Figure S10: Structure prediction and comparison between human and *Plasmodium* subunits.** **A**) Alp5a/Arp3. **B**) Alp5b/Arp2. **C**) PbARPC1/HsARPC1A. **D**) PbARPC2/HsARPC2. **E**) PbARPC4/HsARPC4. Shown are structural prediction of *P. berghei* (first column) and human (second column) Arp2/3 subunits as well as the superimposition of the two structures (third column). Z scores of DALI structural comparisons are indicated.

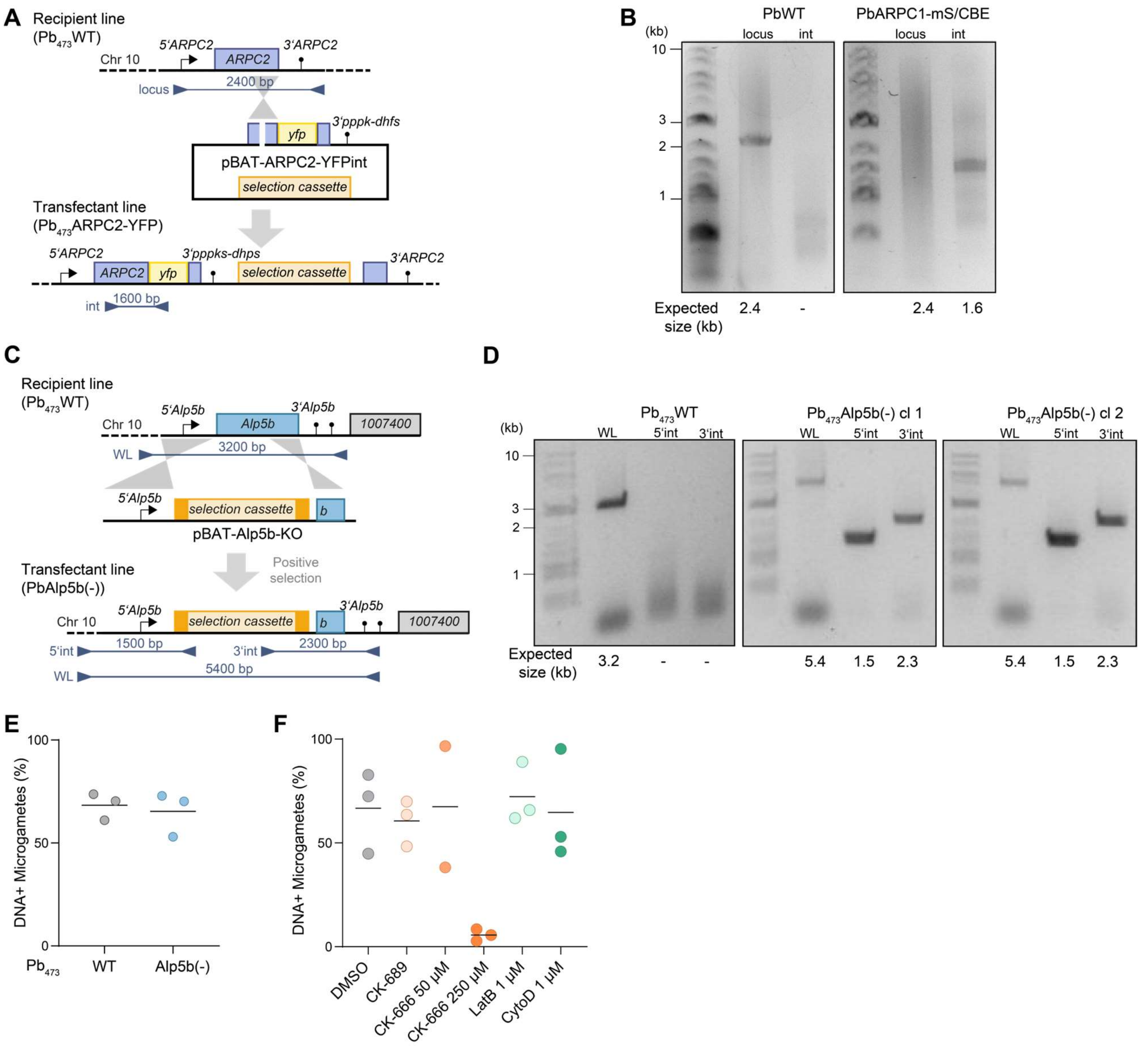


**Figure S11: Generation of parasite lines for localizing and deleting Arp2/3 subunits. A, C**) Scheme of genetic strategy to generate **A**) PbARPC2-YFP and **C**) $Pb_{473}$Alp5b(-). Primers (triangles) and expected fragment sizes used for genotyping (see B, D) are indicated. Not drawn to scale. **B, D**) Genotyping PCR of **B**) PbARPC2-YFP and **D**) $Pb_{473}$Alp5b(-). Primers are indicated in A and C, respectively. **E**) Proportion of microgametes with detectable DNA signal in $Pb_{473}$Alp5b(-). Line at median. **F**) Proportion of microgametes with detectable DNA signal in microgametes after drug treatment. Line at median.

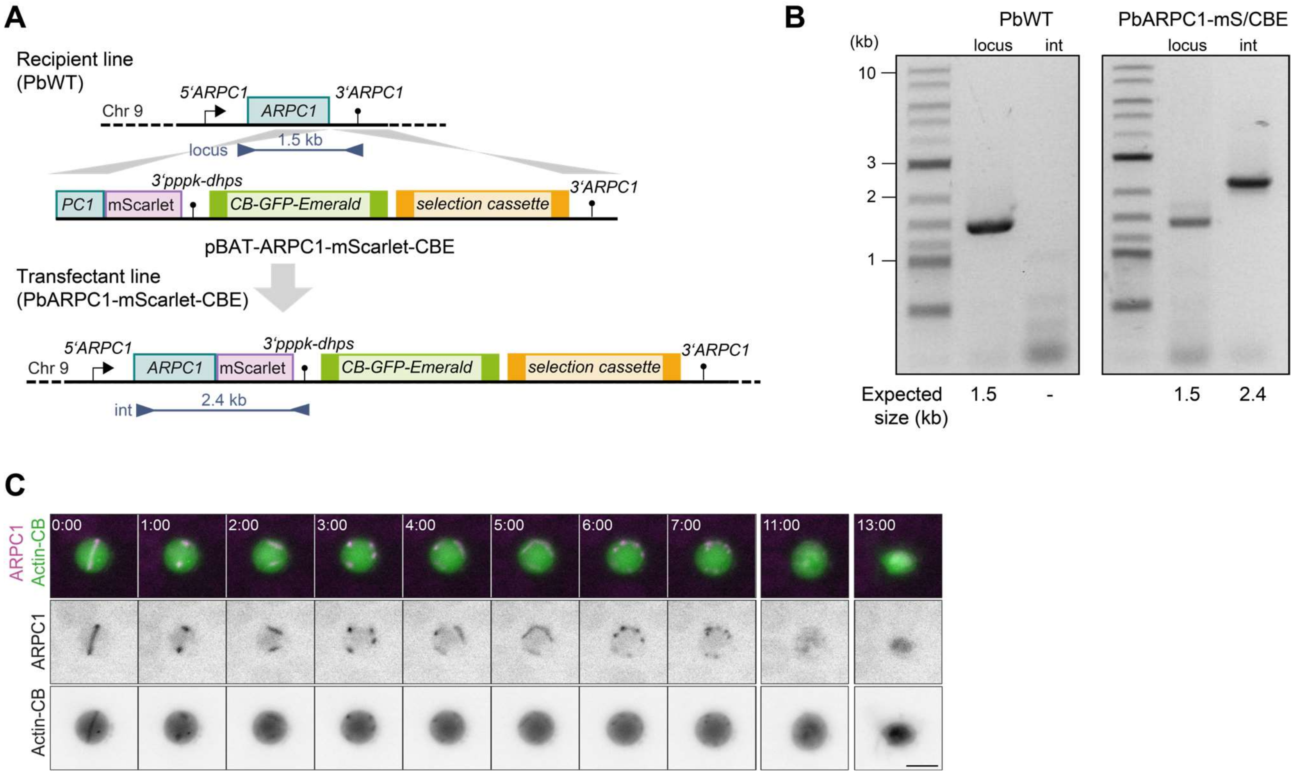


**Figure S12: Generation of parasite lines for knocking out Arp2/3 subunits. A**) Scheme of genetic strategy to generate PbARC40-mScarlet-CBE. Primers (triangles) and expected fragment sizes used for genotyping (see B) are indicated. Not drawn to scale. **B**) Genotyping PCR of PbARC40-mScarlet-CBE. Primers are indicated in A. **C**) Live cell imaging of activated PbARC40-mScarlet-CBE gametocyte. Time points in minutes after start of the movie indicated in upper left corner. Scale bar, 5 µm.

**Table S1: Summary of sporozoite numbers and by bite infections.**

| Parasite line | Oocyst prevalence | Midgut spz #spz/mosquito (#mosquitoes) | Salivary gland spz #spz/mosquito (#mosquitoes) | By-bite infections positive/total mice (mouse strain) |
|---|---|---|---|---|
| **Pb$_{820}$WT** | n.d. | n.d. | 16 000 (20) | 3/3 (TO) |
| **Pb$_{820}$ARC40(-) cl1** | n.d. | n.d. | 0 (20) | 0/3 (TO) |
| **Pb$_{820}$ARC40(-) cl2** | n.d. | n.d. | 0 (20) | 0/3 (TO) |
| **Pb$_{473}$WT** | 100 % | n.d. | 5 700 (10) | 2/2 (TO) |
| **Pb$_{473}$ARC40(-) cl2** | 67 % | n.d. | 0 (10) | 0/2 (TO) |
| **Pb$_{473}$WT** | 83 % | n.d. | 10 000 (10) | n.d. |
| **Pb$_{473}$WT** | 75 % | n.d. | 18 000 (10) | n.d. |
| **Pb$_{473}$ARC40(-) cl2** | 90 % | n.d. | 0 (10) | n.d. |
| **Pb$_{473}$WT** | 47 % | 17 000 (12) | 15 000 (30) | 3/3 (C57Bl/6) |
| **Pb$_{473}$ARC40(-) cl1** | 75 % | 0 (16) | 0 (29) | 0/3 (C57Bl/6) |
| **Pb$_{473}$ARC40(-) cl2** | 67 % | 0 (12) | 0 (30) | 0/3 (C57Bl/6) |
| **Pb$_{473}$WT** | 64 % | 36 000 (14) | 30 000 (15) | n.d. |
| **Pb$_{473}$ARC40(-) cl1** | 77 % | 0 (14) | 0 (20) | n.d. |
| **Pb$_{473}$ARC40(-) cl2** | 55 % | 0 (20) | 0 (15) | n.d. |
| **Pb$_{473}$WT** | 93 % | n.d. | 38 000 (15) | n.d. |
| **Pb$_{473}$ARC40(-)ns** | 83 % | n.d. | 0 (24) | 0/3 (C57Bl/6) |
| **Pb$_{473}$ARC40(-)HAc** | 63 % | n.d. | 27 000 (16) | 3/3 (C57Bl/6) |
| **Pb$_{473}$WT** | 38 % | n.d. | 19 000 (16) | n.d. |
| **Pb$_{473}$ARC40(-)ns** | 26 % | n.d. | 0 | n.d. |
| **Pb$_{473}$ARC40(-)HAc** | 42 % | n.d. | 10 000 (20) | n.d. |

**Table S3: Primers used in this study**

| Primer number | Primer sequence (5‘ → 3‘) |
|---|---|
| P1 | TTCTCCTTTACTCATGGATCCTGCTGCCGCTGCTGCCGCTAAATTATAATTCCAAATAACAACA TTAAAATCG |
| P2 | TTCAATTTCGATATCGAATTATGCCAACACATTCGCAATTGG |
| P3 | GCGGCGGCCGTCTAGAATGCCAACACATTCGCAATTGG |
| P4 | CCATGTTAACACTAGTTAAATTATAATTCCAAATAACAAC |
| P5 | GCGTGGGCCCCCTAGGTGGCTAGCTACGATGTATTT |
| P6 | TACCATCGATAAGCTTGCATGAGTATTATATTGTTACATTT |
| P7 | GGAATTATAATTTAACTAGTGTTAACATGGGATCTGGATCTGGTGGTGGTGGAACCGGTTACC CTTACGATGTTCCTGACTATGCGGGCTATCCCTATGACGTCCC |
| P8 | ATCCTCCAGACAGGCCCACGTGGCCAGCGTAATCTGGAACGTCGTAAGGGTAGCCCATGGC ATAGTCCGGGACGTCATAGGGATAGCCCGCATAGTCAGG |
| P9 | GGAATTATAATTTAACTAGTGTTAACATGG |
| P10 | ATCCTCCAGACAGGCCCAC |
| P11 | CGCCCGCGGCGGCCGTCTAGTTGGCGTGCAATTTGAAAC |
| P12 | CTCCTTTACTCACAGACATACCGGTTCCACCACCACCAGATCCAGATCCTAAATTATAATTCCA AATAACAACATTAAAATCG |
| P13 | ACTATAGGGGACGATCGGTCGTGCACTGCCGAATCTAATTTACATGTATCAAACAATAAGGTA CTATTTG |
| P14 | TTTAGCTGAAATATATTCTCCACCTCCACCAAGATTTTTTTCGGTATTTTCCTCAACATTTTC |
| P15 | AAGGGAGGTGGATCAGCAAGTGGATCTGGTGGAGAATATATTTCAACCTATAAACAATTACAA ACAC |
| P16 | TTCTCCTCCAGATCCTCCAGACAGGCCCACTTAACCACTTCGTCTTTGCTTAATTCGATATAG |
| P17 | AATCTTGGTGGAGGTGGAGAATATATTTCAGCTAAAATTGGAAGTGGAGGACGGATGGTG |
| P18 | TCCACCAGATCCACTTGCTGATCCACCTCCCTTGTACAGCTCGTCCATGCCGAGAGTG |
| P19 | GGAATTATAATTTAACTAGTGTTAACATGGGATCTGGATCTGGTGGTGGTGGAACCGGTATGG TGAGTAAAGGTGAAGCA |
| P20 | ATCCTCCAGACAGGCCCACGTGGCCTTTATATAATTCATCCATTCCACCTGT |
| P21 | AAGCACCAGATCCGGAGCTCGGATAAATAGGGATATGATAAAAT |
| P22 | TTTCCTTCAATTTCGAGCTCATCATATTTGTAATGATGCTTTTTC |
| P23 | ACGATCGGTCGTGCACTGCCGCGGCCGCATTAAAAATTATATGTACCGTTAGTTGG |
| P24 | CACAAATGATGTTTTTTCCTTCAATTTCGTTTTCTTAGCAAAAAATGGGCAG |
| P25 | TAAGTAAGAAAAAACGCGTGGGCCCCCATATTAAGCTCTCATCGTTG |
| P26 | CGATCGGTCGTGCACGACGTGCGGCCGCGTTTATGTAGTGTCTCATTCGTTGGATTG |
| P27 | GAAGAATTACCAACATGCGTTTGCT |
| P28 | CATATAAATACATCGTAGCTAGCC |
| P29 | TTAACATCACCATCTAATTCAACAAG |
| P30 | GTGCATGCTTATGCGCAAAG |
| P31 | TGTGGGTGCCAGCAAACGCA |
| P32 | TCATGCACACACATATGCACA |

| | |
|---|---|
| P33 | GCGATGCGCTTCGTAGTTTGT |
| P34 | CTTTGGTGACAGATACTAC |
| P35 | GTTTCCATATTTATAGATTTGG |
| P36 | CCGCCTACTGCGACTATAGA |
| P37 | GCTCATAATAATTAATGGGATACAAATTCC |
| P38 | TCACAACTTTTCTTCCAGCC |
| P39 | AAAAAATGTGTGAAGTGTTGTAC |
| P40 | CTGCTGCCCGACAACCACTA |
| P41 | GGATATACTTACTCATCATGCCTC |
| P42 | CCTATGCATGTTATTTTCTTGTTCTCTAC |
| P43 | GACTTTGGTGACAGATACTAC |
| P44 | TTCTACTGAAGAGGTTGTGGTC |
